# A validated quantitative method for the assessment of neuroprotective barrier impairment in neurodegenerative disease models

**DOI:** 10.1101/2020.03.06.979930

**Authors:** Vinod Kumar, John D. Lee, Elizabeth J. Coulson, Trent M. Woodruff

## Abstract

The blood brain barrier (BBB) and blood spinal cord barrier (BSCB) are highly specialised structures that limit molecule entry from the blood and maintain homeostasis within the central nervous system (CNS). BBB and BSCB breakdown are associated with multiple neurodegenerative diseases. Given the key role of neuroprotective barrier impairment in neurodegeneration, it is important to identify an effective quantitative method to assess barrier integrity in animal models. In the present study, we developed and validated a quantitative method for assessing BBB and BSCB integrity using sodium fluorescein, a compound that outperformed other fluorescent dyes. We demonstrated using this method that multiple CNS regions progressively increase in permeability in models of Huntington’s disease and amyotrophic lateral sclerosis, while biphasic disruption occurred in a mouse model of Alzheimer’s disease with disease progression. Collectively, we report a quantitative fluorometric marker with validated reproducible experimental methods, that allows the effective assessment of BBB and BSCB integrity in animal models. This method could be useful to further the understanding of the contribution of these neuroprotective barriers to neurodegeneration processes.

## Introduction

The blood brain barrier (BBB) and blood spinal cord barrier (BSCB) are highly specialised epithelial structures linked through intercellular junctional complexes, termed tight junctions. These structures maintain homeostasis within the central nervous system (CNS) by ensuring strict regulation of molecule transport between circulating blood and CNS tissues (Zhao *et al.* 2015). The primary physiological function of these barriers is to maintain the chemical composition of the CNS, and protect against toxic molecules, to support neuronal cell health. BBB and BSCB breakdown, due to disruption of tight junctions or epithelial cell integrity, can lead to altered transport of molecules between the CNS and blood, which can contribute to the pathological processes resulting in progressive structural and functional loss of neurons in multiple neurodegenerative diseases (Ellrichmann *et al.* 2013; Harry & Kraft 2008; Hirsch & Hunot 2009; Lobsiger & Cleveland 2007).

There is evidence to support the notion that neuroprotective barrier impairment occurs within the progression of these neurodegenerative diseases, including in Alzheimer’s disease (AD) (Zlokovic 2011), Parkinson’s disease (PD) (Brochard *et al.* 2009; Carvey *et al.* 2005; Chen *et al.* 2008; Sarkar *et al.* 2014), Huntington’s disease (HD) (Drouin-Ouellet *et al.* 2015) and amyotrophic lateral sclerosis (ALS) (Garbuzova-Davis *et al.* 2007b). Damaged BBB and BSCB integrity and resulting increased permeability can lead to oedema, neurotoxic molecule accumulation, as well as enhancing the infiltration of immune cells to the CNS, which can accelerate the process of neurodegeneration (Arima *et al.* 2013; Bell *et al.* 2010; Stokum *et al.* 2016). This interaction between neuroinflammation, neurodegeneration and neuroprotective barrier integrity can also influence drug bioavailability to the CNS, which may affect the activity of neuroprotective therapies. For example, Tarenflurbil used in AD was shown to have poor BBB penetration, which may have led to its failure in clinical trials (Wan *et al.* 2009).

Given the key involvement of neuroprotective barrier impairment in neurological and neurodegenerative conditions, it is therefore important to identify an effective quantitative method to assess barrier impairment and permeability changes during disease progression in animal models of these conditions.

Previous studies have utilized different dyes to measure the permeability of BBB and BSCB, with Evans blue dye being the most commonly reported (Saunders *et al.* 2015). However, numerous studies have indicated that Evans blue is a substandard marker for BBB permeability, due to significant binding to tissues (Dallal & Chang 1994; Lindner & Heinle 1982), solution instability, *in vivo* toxicity (demonstrated in rats, monkeys, dogs, cats and rabbits), and lack of a validated quantitative determination method. Furthermore, Evans blue relies on binding to the large plasma protein albumin, and subsequent transport across the BBB, which only occurs during later stages of barrier disruption. In contrast to Evans blue, more recent fluorescent-based dyes have been used extensively as a marker for permeability studies including sodium fluorescein (Na-Fl) (Hoffman & Olszewski 1961; Malmgren & Olsson 1980), fluorescein isothiocyanate (FITC) labelled albumin (Boutoille *et al.* 2009; King-VanVlack *et al.* 2003), and FITC labelled dextran-70 (Hoffmann *et al.* 2011; Oliver *et al.* 1984). However, no study to date has directly compared the relative utility of these dyes in assessing BBB and BSCB impairment and provided a robust validated quantitative method.

In the present study, we therefore initially examined these three different dyes (Na-Fl, FITC labelled albumin and FITC labelled dextrans-70) as potential markers for neuroprotective permeability in the brains of lipopolysaccharide (LPS)-injected mice to determine the most suitable and sensitive marker for BBB and BSCB permeability studies. We developed a quantitative method that demonstrated Na-Fl as the most sensitive marker for measuring neuroprotective barrier integrity. We then utilized this method in mouse models of AD, HD, and ALS, where we identified perturbed barrier integrity in different regions of the brain and spinal cord of these animals. Our data therefore reveal a novel quantitative method for monitoring BBB and BSCB integrity in animal models of neurodegeneration.

## Material and methods

### Animals

All animal procedures were approved by the University of Queensland Animal Ethics Committee (Permit Number: 291/15) and conducted in accordance with the Australian Code of Practice for the Care and Use of Animals for Scientific Purposes (8th edition, 2013). All experiments were conducted in compliance with ARRIVE guidelines. This study was not pre-registered, with no randomisation nor blinding performed. Animals were group housed 3 ∼ 4 per cage on arrival in the University of Queensland animal facility. Mice were housed in a 12-h light: dark phase (on at 06:00 h and off at 18:00 h) with room temperature maintained at 24 ± 2 °C. Animals had free access to *ad-libitum* food and water. All experimental procedures were conducted between 10:00 h and 12:00 h. Animal sources and details are outlined below.

#### Wild-type mice

C57BL/6J male mice aged 10-12 weeks old were obtained from the Animal Resources Centre in Canning Vale, Australia. We used a total of 82 mice for selecting and assessing appropriate fluorescent markers by comparing uptake results of the markers FITC dextran, FITC albumin and Na-Fl in the brain and serum of mice over time after an intraperitoneal LPS injection.

#### Transgenic R6/1 mice (transgenic mouse model for Huntington’s disease)

Transgenic R6/1 mice (B6-CBA-Tg (HDexon1) 61Gpb/1J) expressing a transgene containing the human HD gene were obtained from Jackson Laboratory (RRID: IMSR_JAX:002809; (Mangiarini *et al.* 1996)) and bred with female mice on CBA/BL6 background to produce R6/1 transgenic and non-transgenic (wild-type; WT) mice. Male transgenic R6/1 and matched WT littermate mice were used at various stages of disease progression (6 weeks, 12 weeks, 20 weeks and 32 weeks) (Naver *et al.* 2003). We used a total of 40 mice (20 WT and 20 R6/1) to assess BBB/BSCB integrity at four different stages of disease progression.

#### Transgenic hSOD1^G93A^ mice (transgenic mouse model for amyotrophic lateral sclerosis)

Transgenic hSOD1^G93A^ mice (B6-Cg-Tg (SOD1-G93A) 1Gur/J) carrying a mutant form of the human *SOD1* gene were obtained from Jackson Laboratory (RRID: IMSR_JAX:004435; (Rosen *et al.* 1993)) and bred on C57BL/6J background to produce transgenic hSOD1^G93A^ and WT littermates as control mice. Male hSOD1^G93A^ and WT mice were used at various stages of disease progression (30 days, 70 days, 130 days and 150 – 175 days) as described previously (Lee *et al.* 2013). We used a total of 40 mice (20 WT and 20 hSOD1^G93A^) to assess BBB/BSCB integrity at four different stages of disease progression.

#### Transgenic APP/PS1 mice (transgenic mouse model for Alzheimer’s disease)

Double transgenic APP/PS1 mice (B6-Cg-Tg (APPswe, PSEN1dE9)85Dbo/Mmjax) expressing a chimeric mouse/human amyloid precursor protein and a mutant human presenilin 1 were source from Jackson Laboratory (RRID: MMRRC_034832 – JAX; (Jankowsky *et al.* 2004)) and bred on C57BL/6J background to produce transgenic APP/PS1 and WT littermates. Male APP/PS1 and WT mice were used at 4 and 18 months, ages correlating to early, and late stage disease phenotypes (Radde *et al.* 2006; Serneels *et al.* 2009; Rupp *et al.* 2011). We used a total of 20 mice (10 WT and 10 APP/PS1) to assess BBB/BSCB integrity at early and late stages of disease.

### Experimental conditions for quantification and selection of fluorescent marker permeability studies

Fluorescence stability of markers during the extraction and analysis process were first tested by studying effect of different solvents as a loading solution, the effect of solvents during the extraction process, extraction efficiency of the process, the influence of matrix/solvents on the fluorescence of markers, and loss of fluorescence during the extraction and analysis steps (refer to Supplementary methods for full methodology and results). From these validation steps, TRIS buffer (pH 7.4) was identified as the most suitable solvent.

The selection of an appropriate fluorescent marker was accomplished by comparing uptake results of the markers in the brain and serum of mice after an intraperitoneal LPS injection (Clone 0111:B4; Sigma, Cat#L2630, 4 mg kg^-1^ of mice body weight). After 24 hours, mice were then anaesthetized with intraperitoneally injected zolazapam (50 mg kg^-1^) and xylazine (12 mg kg^-1^) to minimize animal suffering during experiments. This was followed by 100 µl intravenous injection of FITC albumin (Sigma, Cat#A9771), FITC dextran (Sigma, Cat#FD40S) and Na-Fl (Sigma, Cat#46960) at 100 mg mL^-1^. After one hour, blood samples were collected via cardiac puncture and allowed to clot at room temperature for 15 minutes. Samples were then centrifuged at 2,000 x *g* for 10 minutes at 4°C for serum collection. After blood collection, mice were immediately perfused transcardially with 50 mL of 1 x PBS to remove any remaining fluorescent marker in the circulation and euthanized via cervical dislocation. Whole brain (which also consists of extracellular fluids such as the cerebrospinal fluid (CSF)) was carefully collected, washed with 1 x PBS, dried with Whatman® gel blotting paper and homogenized in an equal volume of TRIS buffer (pH 7.4). Serum samples were deproteinized with acetonitrile (1:3; Merck, Cat# 1000292500), vortexed, sonicated and centrifuged at 13,000 x *g* for 15 minutes at room temperature. Supernatant samples were evaporated to a dry state using a centrivap concentrator (Labconco) at room temperature and reconstituted in an equal volume of TRIS buffer (pH 7.4). Brain tissue samples were homogenized with milli Q water (1:2 ratio to the tissue weight). Homogenized brain samples were deproteinized with acetonitrile (1:6 ratio to the tissue weight) and processed via a similar method used for serum samples. Dried brain samples were reconstituted in equal weight-volume of TRIS buffer (pH 7.4). 10 µl of reconstituted solutions were diluted with 200 µl of TRIS buffer (pH 7.4) in 96-well black plate (Corning Inc, Cat#3603) along with standard curve samples in same matrix. The fluorescence levels were measured using a microplate reader (FlexStation 3) with λ_ex_ 460 nm and λ_em_ 515 nm for Na-Fl, λ_ex_ 495 nm and λ_em_ 520 nm for FITC-albumin and λ_ex_ 490 nm and λ_em_ 530 nm for FITC-dextran respectively. The amount of fluorescent marker was calculated from fluorescence levels via a standard curve prepared in a similar manner using brain and serum obtained from mice without any treatment. Raw fluorescent data values obtained for serum and brain are located at Open Science Framework (DOI 10.17605/OSF.IO/Q6XGB). Fluorescence uptake was calculated by comparing ratio of fluorescent marker level in the brain with the level of fluorescent marker in the serum. Fluorescence values were corrected with the fluorescence of blank brain and serum samples obtained from mice without any fluorescent marker or LPS treatment and treated with the same method as above.

### Application of a validated quantitative fluorometric method in mouse models of neurodegeneration

The applicability of the developed and validated quantitative fluorometric method was assessed by performing neuroprotective barrier integrity studies in various chronic mouse models of neurodegeneration. Mice (*n* = 5) at various disease stages (Table 1) were anaesthetized with an intraperitoneal injection of zolazapam (50 mg kg^-1^) and xylazine (12 mg kg^-1^) to minimize animal suffering during the experiments. Anaesthetized mice were then intravenously administered with 100 µl of Na-Fl (100 mg mL^-1^) via tail vein. After 15 minutes from injection, blood samples were collected by cardiac puncture for serum extraction. Animals were transcardially perfused immediately with 1 x PBS to remove circulating Na-Fl and euthanized via cervical dislocation. Following this, whole brain and spinal cord tissue was collected and different brain regions (cortex, striatum, hippocampus and cerebellum) were isolated on ice using a mouse coronal brain matrix (Cat#U040-C, ProSciTech) under the microscope as per the suggested protocol (Spijker 2011). Samples were stored at −80°C for further processing and analysis using the developed quantitative fluorometric method as described above. Fluorescence levels in brain and spinal cord samples were measured as describe above. Fluorescence values were back calculated for the level of Na-Fl in the serum and tissues using the standard curve samples prepared in the same matrix and analyzed as above. Results are presented as relative levels of fluorescence uptake in R6/1, hSOD1^G93A^ and APP/PS1 mice compared with expression in their respective litter-matched non-transgenic WT controls. Complete data values obtained prior to normalisation to WT controls are also found at Open Science Framework (DOI 10.17605/OSF.IO/Q6XGB).

**Table 1:**
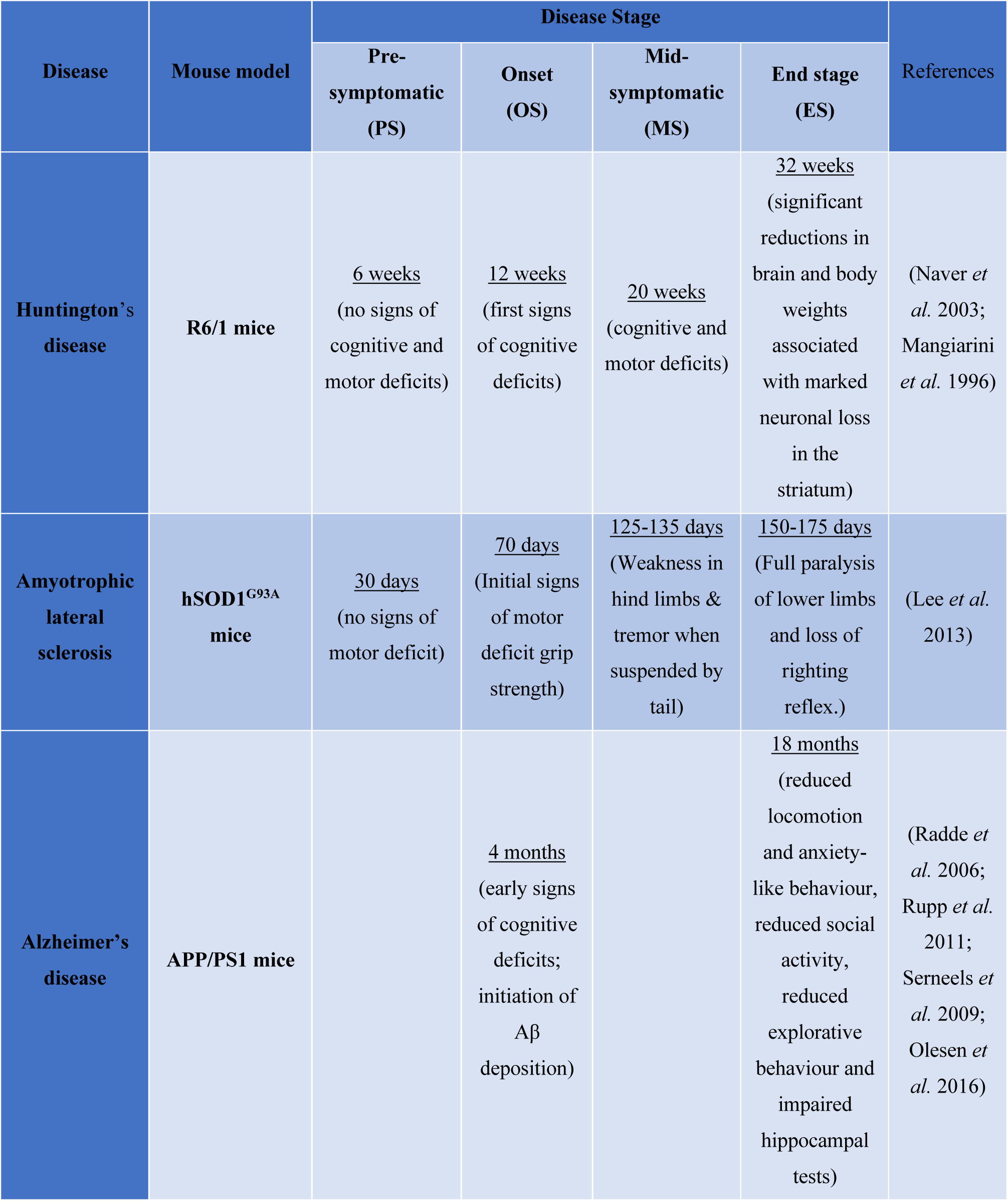
Different stages of neurodegeneration in transgenic mouse models used in this study.

## Statistical analysis

All results are reported as mean ± standard error of mean and statistical analysis performed using GraphPad Prism 8.3.1 (RRID:SCR_002798). No sample size calculation was performed prior to starting the experiments. No tests for outliers (i.e. no exclusion of data points) and no exclusion criteria were pre-determined for the study. Normality was tested using Shapiro – Wilk test. The statistical difference between FITC albumin, FITC dextran and Na-Fl for percentage uptake in the brain were analyzed using a one-way ANOVA and a *post-hoc* Tukey’s test. The statistical difference for levels of fluorescence uptake between WT and R6/1, WT and hSOD1^G93A^ and WT and APP/PS1 mice were analyzed using a two-tailed Student’s t-test for each stage of disease progression. The differences were considered significant when *P* < 0.05.

## Results

### Sodium fluorescein has greater sensitivity as an indicator of BBB disruption than FITC-dextran or FITC-albumin

Selection of an appropriate fluorescent marker for permeability studies was based upon the percentage uptake of the fluorescent marker, which was quantitatively determined via a validated fluorometric method. We utilized peripheral administration of LPS to induce BBB disruption after 24 hours as previously documented (Banks *et al.* 2015; Nonaka *et al.* 2005). We then determined the uptake of fluorescent dye into the brain 1 hour after intravenous dye injection. The percentage uptake of Na-Fl (∼50%) was found to be the highest compared to either FITC-dextran-70 (∼1%) or FITC-albumin (∼17%) (**Figure 1A**). We next extended this analysis by comparing BBB permeability changes using FITC-albumin and Na-Fl (both having detectible uptake in the brain after 1 hour), over 24 hours. Fluorescence uptake of FITC-albumin in the brain was significantly increased at 0.25 hour (1.4% vs. 4.8%), 1 hour (6.8% vs. 16.7%) and 24 hours (8.4% vs. 24%) in saline-injected versus LPS-injected mice respectively (**Figure 1B**). Fluorescence uptake of Na-Fl in the brain was also significantly increased at 0.25 hour (8.5% vs. 40%), 1 hour (10.4% vs. 67.7%) and 24 hours (3.8% vs. 17.2%) in saline-injected versus LPS-injected mice respectively (**Figure 1C**). Notably, the fold-increases in dye uptake between saline and LPS groups, were significantly larger for Na-Fl than FITC-albumin, peaking 1 hour after dye injection. We therefore identified 1-hour Na-Fl to be an ideal method to progress for future validation.

**Figure 1.**
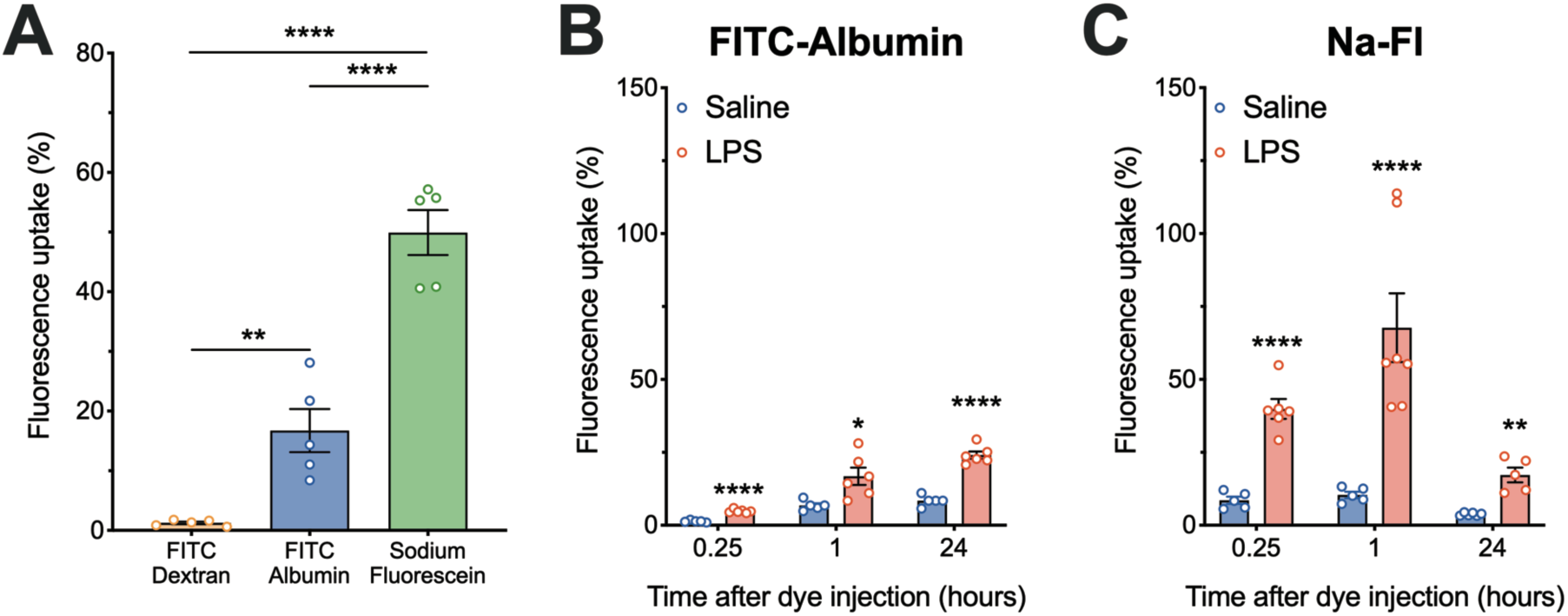
Brain fluorescence uptake profile of three different fluorescent markers. **(A)** Fluorescence uptake of sodium fluorescein (Na-Fl), FITC-albumin and FITC-dextran in LPS treated mice after 1 hour of *i.v.* injection of fluorescent marker through tail vein (*n* = 5 animals). Fluorescence uptake of Na-Fl **(B)** and FITC-albumin **(C)** in control (blue) and LPS-administered mice (red) after 0.25, 1 and 24 hours of *i.v.* injection of fluorescent marker through tail vein (*n* = 5 – 7 animals). Uptake values are expressed as % ratio of fluorescent marker level in brain to serum. Data are expressed as mean ± SEM. * *P* < 0.05, ** *P* < 0.01, **** *P* < 0.0001, one-way ANOVA.

### Progressive breakdown of neuroprotective barriers in the R6/1 transgenic mouse model of Huntington’s disease

We first applied our method to investigate BBB and BSCB integrity at different disease stages in the R6/1 transgenic mouse model of HD. We observed no change in Na-Fl uptake in cortex, striatum, cerebellum and spinal cord between WT and R6/1 transgenic mice at 6 weeks of age, which is considered to be a pre-symptomatic stage of disease. Na-Fl uptake in the cortex and striatum progressively increased in R6/1 mice at 12 (1.7-fold, 1.8-fold), 20 (2.1-fold, 2.4-fold) and 32 (2.3-fold, 2.9-fold) weeks when compared to WT mice (**Figure 2A** and **2B**). The cerebellum was less affected, with Na-Fl uptake only increased by 2.0-fold in R6/1 mice at 32 weeks of age (**Figure 2C**). Interestingly, Na-Fl was also increased in the spinal cord of R6/1 mice at 20 and 32 weeks by 2.7-fold respectively (**Figure 2D**).

**Figure 2:**
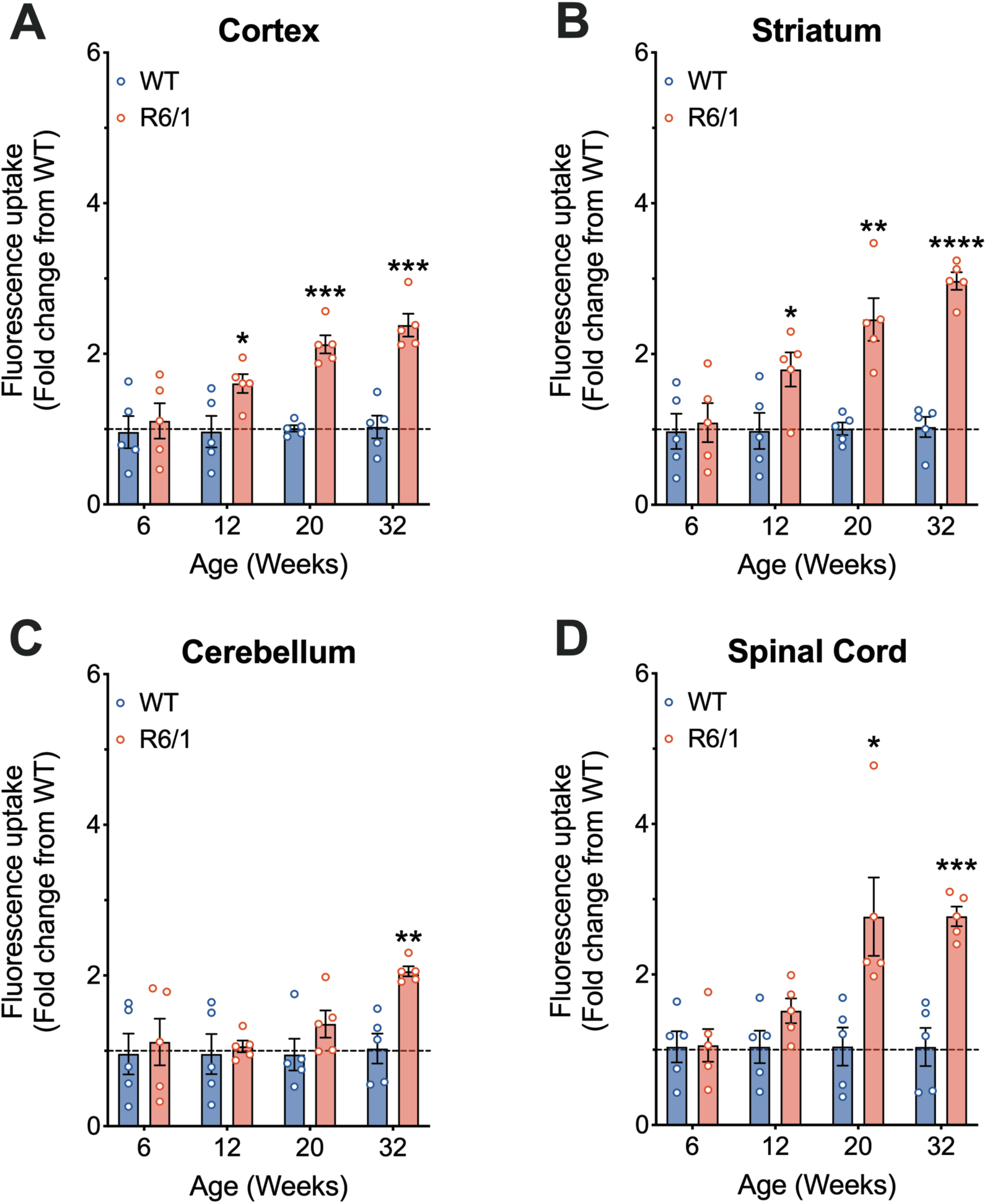
Progressive increase of Na-Fl in the brain and spinal cord of R6/1 Huntington’s mice throughout disease progression. **A** and **B** show increased Na-Fl uptake in cortex and striatum of R6/1 mice at 12, 20 and 32 weeks when compared to wild-type (WT) mice (*n* = 5 animals). **C** shows an increased Na-Fl extravasation in cerebellum of R6/1 mice at 32 weeks when compared to WT mice (*n* = 5 animals). **D** shows increased Na-Fl uptake in the spinal cord of R6/1 mice at 20 and 32 weeks of age when compared to WT mice (*n* = 5 animals). Data are presented as mean ± SEM. * *P* < 0.05, ** *P* < 0.01, *** *P* < 0.001, **** *P* < 0.0001, Student’s t-test for each time point.

### Breakdown of neuroprotective barriers in the hSOD1^G93A^ transgenic mouse model of amyotrophic lateral sclerosis

Next, we assessed neuroprotective barrier integrity in hSOD1^G93A^ transgenic mice. Na-Fl uptake was measured in cortex, striatum, cerebellum and spinal cord at progressive ages of disease progression. Similar to R6/1 mice, we observed no changes in Na-Fl uptake in cortex, striatum, cerebellum and spinal cord between WT and hSOD1^G93A^ transgenic mice at the pre-symptomatic age of 30 days. However, Na-Fl uptake progressively increased in the cortex of hSOD1^G93A^ mice at 70 (1.2-fold), 130 (1.5-fold) and 175 (1.6-fold) days of age (**Figure 3A**). Na-Fl also increased in the striatum of hSOD1^G93A^ mice at these same ages (**Figure 3B**). BBB permeability increases were also observed in the cerebellum of hSOD1^G93A^ mice, with significant Na-Fl uptake observed at 70, 130 and 175 days (**Figure 3C**). Finally, BSCB permeability was also detected, with Na-Fl uptake in the spinal cord gradually increased in hSOD1^G93A^ mice by 1.1-, 1.3- and 2.0-fold at 70, 130 and 175 days respectively (**Figure. 3D**).

**Figure 3:**
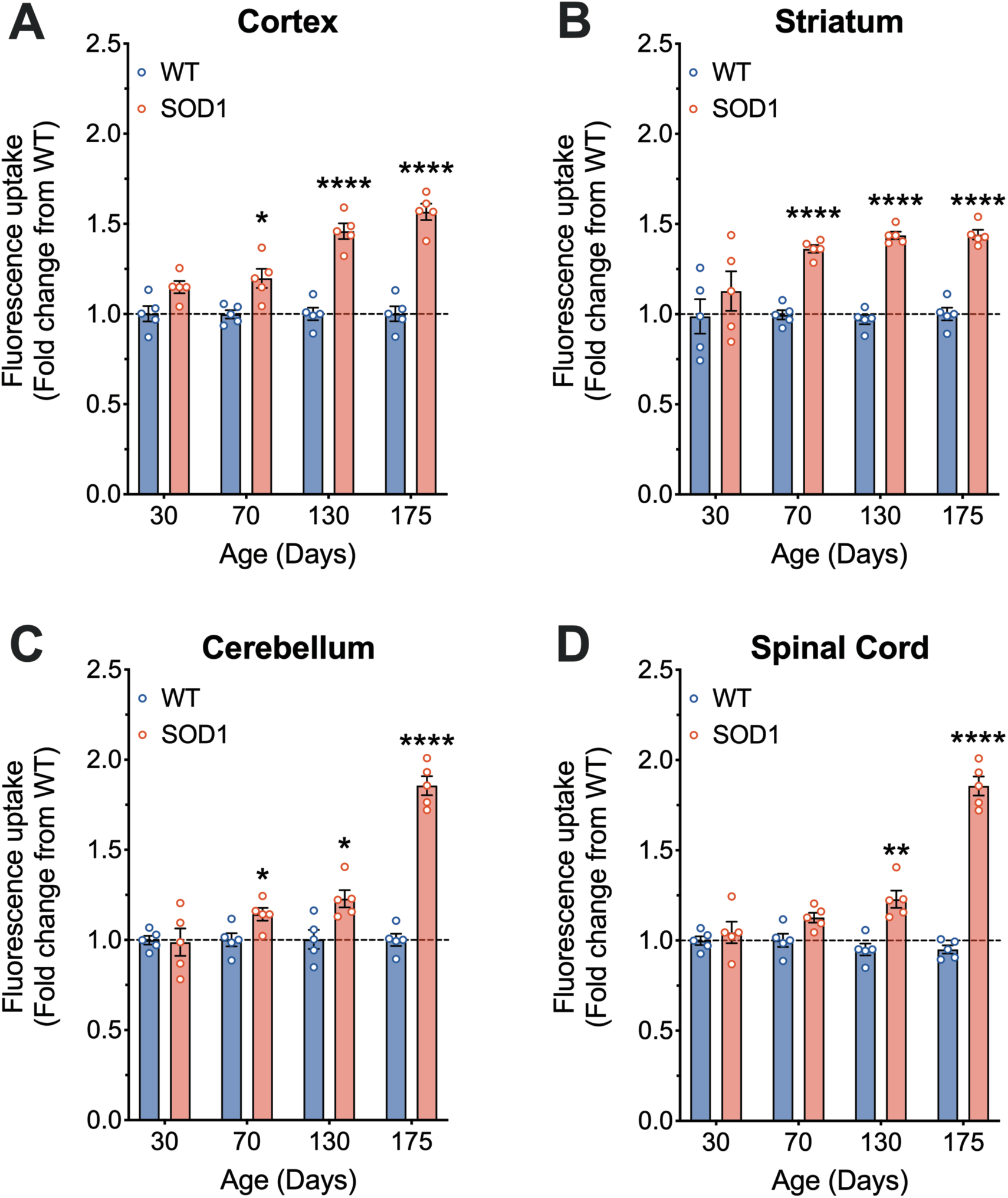
Increased uptake of Na-Fl in the brain and spinal cord of hSOD1^G93A^ amyotrophic lateral sclerosis mice throughout disease progression. **A, B** and **C** shows increased fluorescence uptake of Na-Fl in cortex, striatum and cerebellum of hSOD1^G93A^ (SOD1) mice at 70, 130 and 175 days when compared to wild-type (WT) mice (*n* = 5 animals). **D** shows increased Na-Fl uptake in the spinal cord of SOD1 mice at 130 and 175 days of age when compared to WT mice (*n* = 5 animals). Data are presented as mean ± SEM. * *P* < 0.05, ** *P* < 0.01, **** *P* < 0.0001, Student’s t-test for each time point.

### Neuroprotective barriers in the APP/PS1 transgenic mouse model of Alzheimer’s disease display biphasic alterations at different diseases stages

To measure possible alterations in neuroprotective barriers of APP/PS1 transgenic mice, Na-Fl uptake was measured in cortex, striatum, hippocampus, cerebellum and spinal cord at an early (4 months) and late (18 months) stage of disease. Brain and spinal cord tissues from APP/PS1 transgenic mice at the early stages of disease showed an overall increase in barrier permeability (**Figure 4**), with significant increases in Na-Fl uptake observed in the striatum (**Figure 4C**) and spinal cord (**Figure 4E**). Notably, by 18 months of age, these same tissues had reduced levels of Na-Fl, with decreases of up to 50% observed in brain regions and spinal cord (**Figure 4B**).

**Figure 4:**
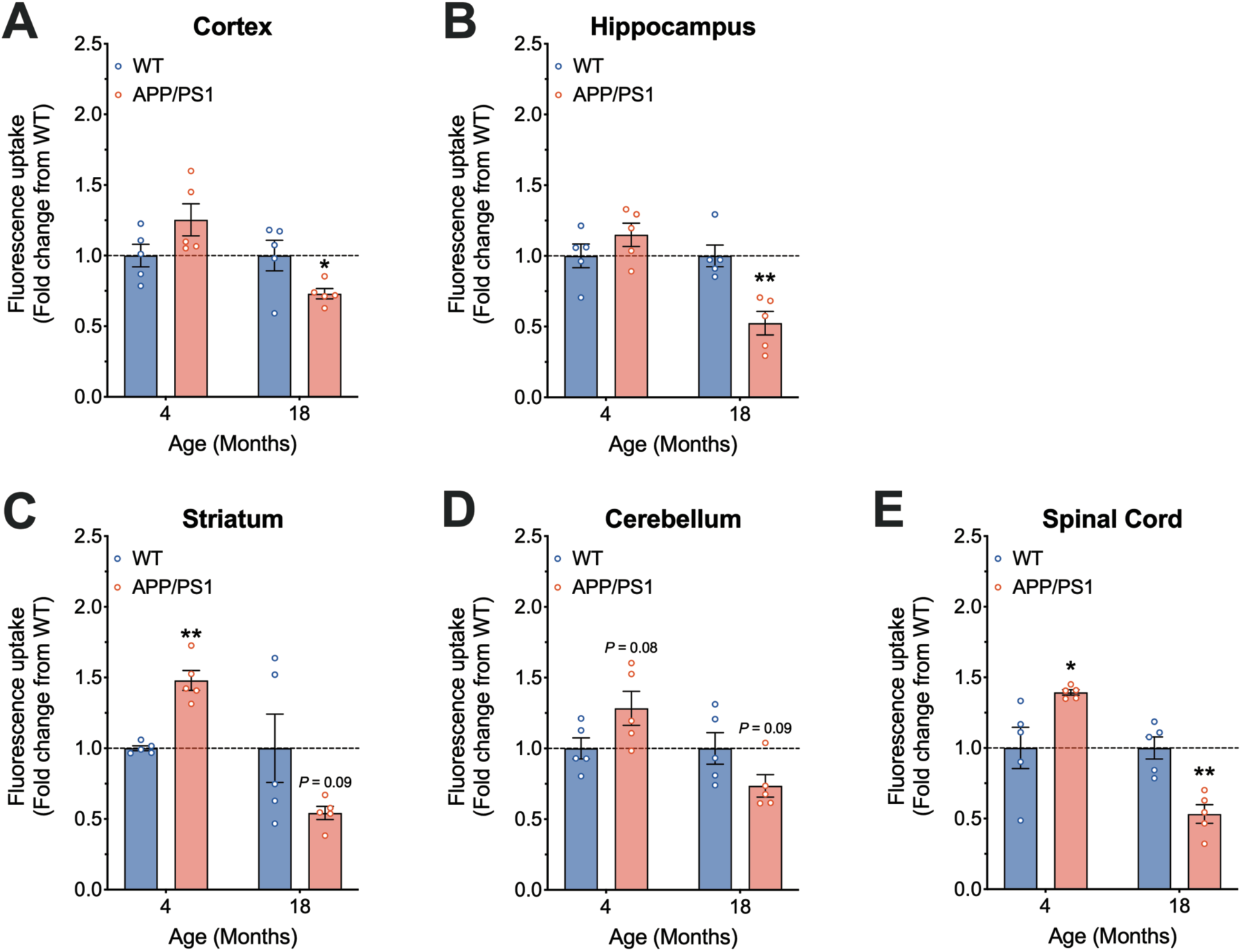
Biphasic brain and spinal cord Na-Fl uptake in the APP/PS1 mouse model of Alzheimer’s disease. Fluorescence was examined in the cortex of APP/PS1 and wild-type (WT) mice at an early (4 months) and late (18 months) age (*n* = 5 animals). Na-Fl uptake was examined in the (**A**) cortex, (**B**) hippocampus, (**C**) striatum, (**D**) cerebellum, and (**E**) spinal cord (*n* = 5 animals). Data are presented as mean ± SEM. * *P* < 0.05, ** *P* < 0.01, Student’s t-test for each time point. *P* values < 0.1 are also identified on the graph.

## Discussion

BBB and BSCB breakdown are associated with a number of neurodegenerative diseases, including AD, PD, HD and ALS (Desai *et al.* 2007; Di Pardo *et al.* 2017; Garbuzova-Davis *et al.* 2007a; Garbuzova-Davis & Sanberg 2014; Garbuzova-Davis *et al.* 2007b; Gray & Woulfe 2015). The understanding of the underlying mechanisms that initiates the opening or closure of the BBB and BSCB is therefore of therapeutic significance in many neurodegenerative diseases and is commonly investigated by researchers. However, methods to investigate BBB and BSCB permeability *in-vivo* are often associated with technical difficulties that depend on subjective quantification of fluorescence images, or semi-quantitative evaluation of endogenous molecules. To address this, prior studies have utilized an abundance of fluorescent markers for *in-vivo* and *in-vitro* permeability studies, each with their own strength and weaknesses (Saunders *et al.* 2015; de Vries *et al.* 1995; Ravenstijn *et al.* 2008; de Lange *et al.* 1998). Methods used for the detection of these fluorescent markers also display great variation in their quantitation methodology between studies. Selection of an ideal validated method for assessing neuroprotective barrier permeability is therefore difficult due to lack of prior published results for quantitative determination. The present research, supported by previously published methods (de Lange *et al.* 1998; Ravenstijn *et al.* 2008; de Vries *et al.* 1995), therefore instigated the selection and identification of a suitable fluorescent marker for CNS permeability studies using a validated fluorometric method of quantitative analysis. The method was developed using an acute LPS-induced BBB breakdown model and then its application verified by assessing neuroprotective barrier permeability in various mouse model of neurodegeneration.

The BBB and BSCB are highly specialized epithelial structures that are important in maintaining the CNS homeostasis and preventing the entry of foreign toxic substances. It undergoes degeneration during disease processes of neurodegenerative diseases and has a re-emerging interest as a mechanistic and therapeutic target for intervention. In spite of their limitations, use of dyes to detect barrier breakdown have remained widespread, where numerous studies have utilized different dyes including, Evans blue, fluorescein, horse radish peroxidase, dextran, albumin and trypan blue (Saunders *et al.* 2015). To the best of our knowledge, our study is the first to compare three widely used fluorescent dye markers (FITC-dextran, FITC-albumin and Na-Fl) to determine the most suitable marker to quantitatively evaluate BBB integrity. We demonstrate that 1-hour after the injection of these dyes to LPS-induced BBB-injured mice, Na-Fl showed the greatest percentage uptake in the brain. Although this has not been directly compared previously, this is consistent with previous studies suggesting that Na-Fl only binds weakly to proteins, and is therefore considered to be an effective low molecular weight marker for BBB studies in contrast to dyes that bind to proteins (Wolman *et al.* 1981). Although Na-Fl cannot be detected by visual examination, a quantitative fluorometric method utilized in the current study may enable detection of subtle alterations in BBB/BSCB permeability and be the earliest and the most sensitive indicator of BBB/BSCB disruption (Kaya & Ahishali 2011). Our studies also demonstrated that Na-Fl uptake in the brain was reduced after 24 hours of injection (when compared to 15 minutes and 1 hour), suggesting rapid clearance of Na-Fl in the brain parenchyma. Collectively, as small molecule, Na-Fl is considered to be less toxic than other dye tracers, is freely diffusible, minimally binds to proteins and can be evaluated quantitatively and reproducibly. We therefore posit this method provides an ideal marker of assessing neuroprotective barrier integrity. In contrast to Na-Fl, FITC-albumin reflects a significant increase in time dependant percentage uptake, supporting its potential as a marker for longer-duration permeability studies.

In pathological conditions, BBB permeability is increased due to various biochemical mediators (Abbott 2000). BBB permeability also increases with ageing (Erdő *et al.* 2017), and our data obtained from WT mice from the transgenic mouse colonies across different ages supports this (Supplementary Figure 1). Increased permeability favours the transport of peripheral immune cells, inflammatory products and other circulating substances to the brain and spinal cord, resulting in breakdown of tight junction molecules and ultimately, disruption of BBB and BSCB (Ballabh *et al.* 2004; Abbott *et al.* 2010; de Vries *et al.* 1997). BBB and BSCB disruption and malfunction are involved in various neurological disorders including AD, PD, HD and ALS (Desai *et al.* 2007; Di Pardo *et al.* 2017; Garbuzova-Davis *et al.* 2007a; Garbuzova-Davis & Sanberg 2014; Garbuzova-Davis *et al.* 2007b; Gray & Woulfe 2015). Using different mouse models of HD, ALS and AD, we investigated barrier integrity in different regions of the brain and spinal cord using our quantitative method of Na-Fl extravasation. We demonstrated that there is an early disruption of BBB and BSCB permeability in R6/1 mouse model of HD as we showed increased Na-Fl uptake in cortex, striatum and spinal cord, in parallel with the cognitive and motor deficits present in this model (Ayton *et al.* 2019; Brooks *et al.* 2012). This early disruption of BBB is further supported by previous studies showing early vascular impairment and BBB dysfunction in R6/2 mice, with a reduction in expression of tight junction proteins claudin-5 and occludin, leading to increased paracellular permeability (Di Pardo *et al.* 2017; Drouin-Ouellet *et al.* 2015; Luissint *et al.* 2012). This validates the applicability of this fluorometric method to measure subtle changes in BBB/BSCB integrity.

We also demonstrated that there is an early disruption of BBB and BSCB permeability in hSOD1^G93A^ mouse model of ALS based on increased Na-Fl uptake in cortex, striatum, cerebellum and spinal cord. Similar to HD, these findings support previous studies demonstrating damage to BSCB integrity in hSOD1^G93A^ mice via ultrastructural capillary alterations and increased leakage in the spinal cord at both early and late stages of disease (Garbuzova-Davis *et al.* 2007a; Garbuzova-Davis *et al.* 2007b; Lee *et al.* 2017; Zhong *et al.* 2008). These findings are further supported by numerous studies demonstrating decreased expression of tight junction proteins integral to the integrity of the BSCB, such as zona occludens-1, occludin and claudin-5 in the spinal cord of hSOD1 transgenic mice and ALS patients (Henkel *et al.* 2009; Meister *et al.* 2015; Winkler *et al.* 2014; Zhong *et al.* 2008). Collectively, our study complements these previous findings given our method was able to detect early barrier impairments very early in disease, at ages that precede significant motor neuron death (Lee *et al.* 2013), which may have pathophysiological relevance. As neuroinflammation and microgliosis are commonly correlated with barrier impairment (Erdő *et al.* 2017), it will be of interest to explore these association in future studies, particularly as neuroinflammation is a key driver of pathology in hSOD1^G93A^ mice (Chiarotto *et al.* 2019).

We also demonstrated an apparent biphasic BBB/BSCB disruption in APP/PS1 mouse model of AD. Unlike HD and MND, neuroprotective barrier disruption in APP/PS1 mouse model of AD was only evident in early in the disease, at an age where there is minimal cognitive dysfunction (Zhu *et al.* 2017; Bien-Ly *et al.* 2015). Interestingly, unlike the other disease models studied, there was a notable decrease in Na-Fl uptake in brain and spinal cord regions at the late stage of disease, an age where is significant cognitive decline (Gengler *et al.* 2010; Milne *et al.* 2019). Although it is possible that the result is due to a faster clearance of the dye from a highly permeable BBB, the decrease in dye uptake in APP/PS1 mice late in disease is consistent with previous studies demonstrating thickening of the basement membrane in AD model mice and humans (Hawkes *et al.* 2013; Lepelletier *et al.* 2017). Although the causes of the basement membrane thickening in AD have not been fully elucidated to date, decreased clearance of amyloid beta or increased astrocyte production of basement membrane proteins like perlecan and fibronectin have been suggested to contribute to this thickening of basement membrane (Deane & Zlokovic 2007; Okoye & Watanabe 1982; Thomsen *et al.* 2017). Importantly, the progressive molecular changes in barrier permeability of multiple neurodegenerative disease models tested in previous studies mirrored the changes in Na-Fl levels in the brain and spinal cord of our current study. This strongly validates the sensitivity and accuracy of our validated fluorometric method to measure subtle changes in BBB/BSCB integrity. This may have implications for future preclinical studies that aim to correlate the extent of barrier impairment and permeability changes with disease progression.

In summary, we have identified a validated, effective quantitative method for assessing BBB and BSCB integrity using Na-Fl. We demonstrated using this method that multiple CNS regions progressively increase in permeability in the R6/1 mouse model of HD and the hSOD1^G93A^ mouse model of ALS. By contrast, whilst increased BBB/BSCB permeability was observed in young APP/PS1 mouse model of AD, later stage diseased mice showed marked reduction in CNS permeability. Collectively, we report a quantitative fluorometric marker with validated reproducible experimental methods, that allows the effective assessment of BBB and BSCB integrity in animal models. This method could be useful to further the understanding of the contribution of these neuroprotective barriers to neurodegeneration processes, and to assess the efficacy of neuroprotective agents.

## Abbreviations used

AD: Alzheimer’s disease
ALS: amyotrophic lateral sclerosis
BBB: blood brain barrier
BSCB: blood spinal cord barrier
CNS: central nervous system
FITC: fluorescein isothiocyanate
HD: Huntington’s disease
LPS: lipopolysaccharide
Na-Fl: sodium fluorescein
PD: Parkinson’s disease
RRID: Research Resource Identifier (see scicrunch.org)
WT: wild-type

## Acknowledgements and conflict of interest disclosure

The research was funded by grants from the MNDRIA to TMW (GIA1865) and JDL (GIA1830), and a National Health and Medical Research Council (NHMRC) Project grant (APP1082271) to TMW. The authors would like to sincerely thank the University of Queensland Biological Resources for the animal care and husbandry. We also thank Maryam Shayegh for her technical support with genotyping mice. The authors declare that they have no competing interests. An earlier version of this manuscript was uploaded to bioRxiv: https://www.biorxiv.org/content/10.1101/2020.03.06.979930v1.

## Author’s contributions

VK, JDL, and TMW conceived the project. VK, JDL, EJC and TMW designed the study. VK and JDL performed experiments. All authors contributed to the analyses and/or interpreted the data. VK and JDL wrote the paper with TMW, with editing contributions from EJC. All authors read and approved the final manuscript.

