## Supplementary for "A validated quantitative method for the assessment of neuroprotective barrier impairment in neurodegenerative disease models"

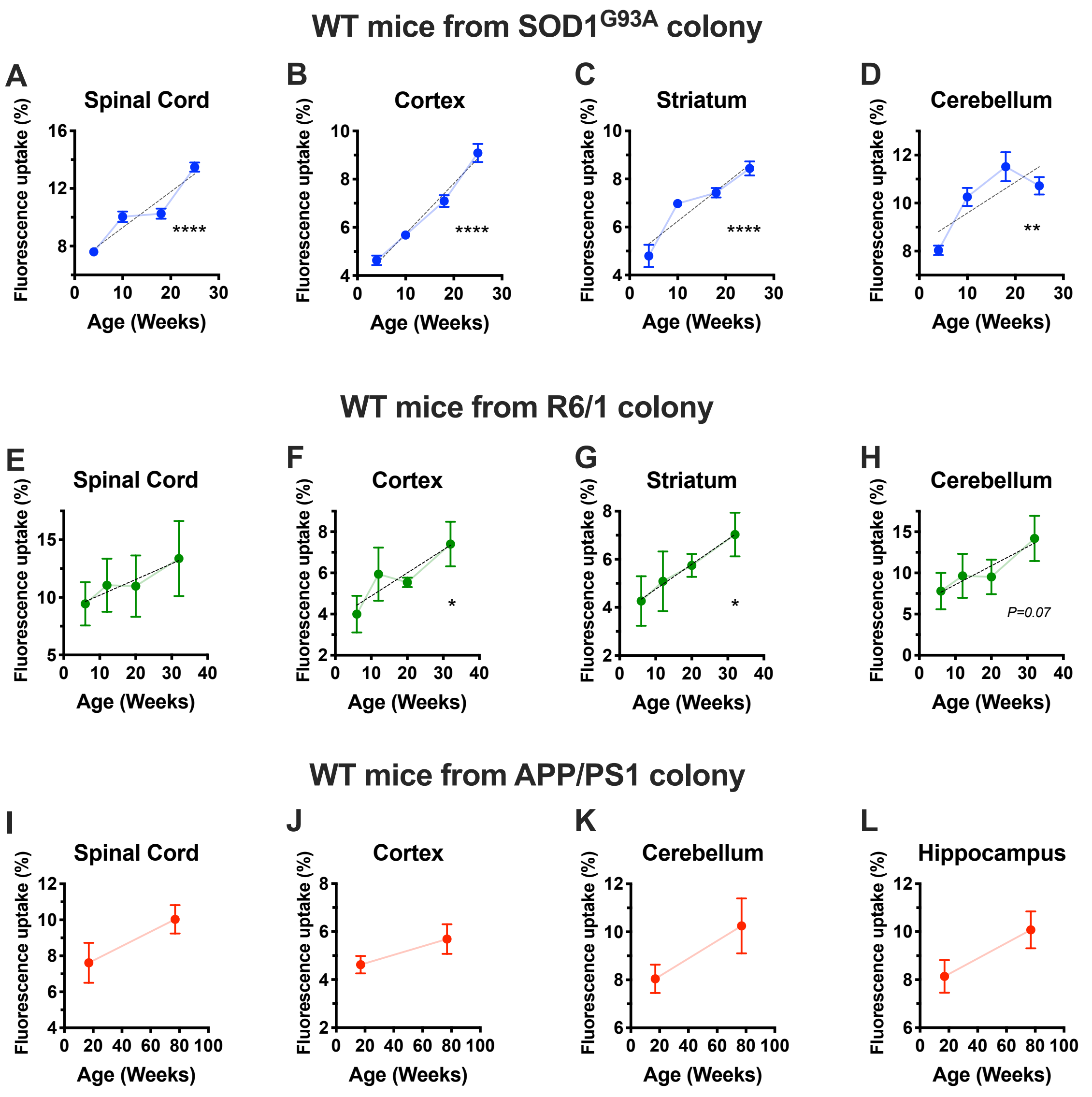


**<u>Supplementary Figure 1</u>: Age-induced increases in neuroprotective barrier permeability to Na-Fl.** Graphs represent % fluorescence uptake of Na-Fl in spinal cord (**A**, **E** and **I**), cortex (**B**, **F** and **J**), striatum (**C**, **G** and **K**) and cerebellum (**D**, **H** and **L**) regions of wild-type mice from SOD1$^{G93A}$ (**A**-**D**), R6/1 (**E**-**H**) and APP/PS1 (**I**-**L**) transgenic colonies. Data expressed as mean ± SEM of n=5 mice per group. Dotted line in each panel represents the linear regression, with significance determined as slope deviation from zero; *P<0.05, **P<0.01, ****P<0.0001.

### SUPPLEMENTARY METHODOLOGY

**Detailed experimental conditions for selection and quantification of fluorescent markers**

A fluorescent marker's fluorescence emission is greatly influenced by its interaction with solvents. Some solvents enhance fluorescence, while others cause a quenching effect on fluorescence. Hence, it was critical to initially identify the extent and influence of solvents on the fluorescence of the selected markers in order to identify best solvent conditions by which fluorescence stability could be maintained during quantification. In the present study, the fluorescence stability of markers during the extraction and analysis process was tested by studying: (i) effect of solvents as a loading solution, (ii) effect of solvents during extraction process, (iii) extraction efficiency of process, (iv) influence of matrix/solvents over fluorescence of markers, and (v) loss of fluorescence or marker during whole extraction and analysis steps.

*Solvent effect as loading solution:* A small volume of final sample (i.e. 10 µl) along with 200 µl of loading solution was added to one well of 96 well clear bottom black plate. The ideal loading solution was selected out of various standard solvents used during extraction and sample preparation. 10 µl of either fluorescent marker (i.e. 0.1% Na-Fl or 1% FITC-albumin or 1% FITC-dextran) along with 200 µl of solvents were auto-mixed for 5 minutes and fluorescence of the resulting solution was determined using a Flexstation fluorescent plate reader. The goal was to identify a loading solution that could be useful during fluorescence determination with minimum effect on fluorescence stability of the fluorescent marker.

**Figure A** illustrates the effect of various solvents on the fluorescence output of the three fluorescent markers tested. Organic solvent 1-butanol enhanced the fluorescence of Na-Fl, FITC-albumin and FITC-dextran. Other solvents such as RIPA buffer, methanol and acetonitrile resulted in fluorescence loss, likely due to chemical interaction of solvent with the markers. Out of all solvents, TRIS buffer at pH 7.4 was identified as the ideal loading solvent for quantitative fluorometric determination, with reproducible results close to the standard solution when analysed alone in the absence of any solvent (no solvent).

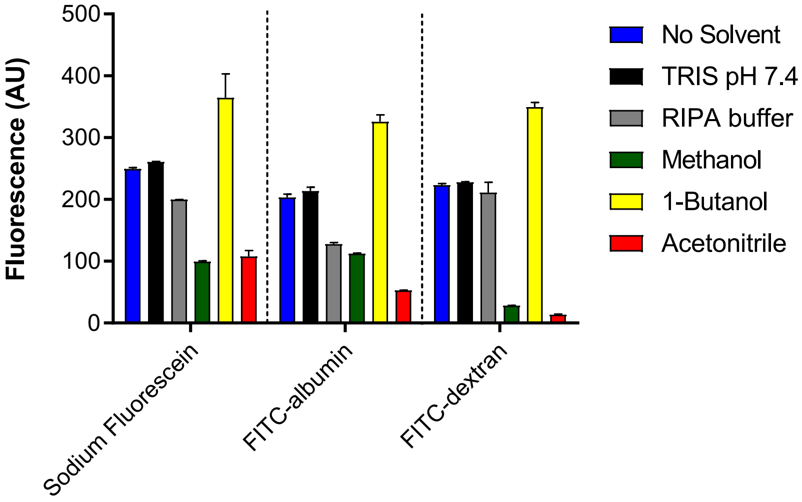


**Figure A: Effect of solvents as loading solution**: Bar graph represents fluorescence level of sodium fluorescein (Na-Fl), FITC-albumin and FITC-dextran in various solvents. Results are compared with fluorescence of 10 µl of fluorescent marker (solution freshly prepared in TRIS buffer pH 7.4) as represented by blue bar graph without any loading solvent. Data represents mean ± SEM of n=4 values.

Extraction solvent effect: The effect of organic solvents on the fluorescence of markers during the extraction procedure was studied next. This could be helpful in selection of the ideal organic solvent capable of efficiently extracting fluorescent markers from tissues and serum samples. To test this, 10 µl of fluorescent marker (i.e. 0.1% Na-Fl or 1% FITC-albumin or 1% FITC-dextran) was mixed and vortexed with 390 µl of organic solvent (i.e. methanol or ACN or 1-butanol) in one batch. In a second batch, 10 µl of either of fluorescent marker (i.e. 0.1% Na-Fl or 1% FITC-albumin or 1% FITC-dextran) was mixed and vortexed with 390 µl of one of the organic solvents (i.e. methanol or ACN or 1-butanol) followed by drying using a centrivap evaporator at room temperature. Dried extract was dissolved in 400 µl of TRIS buffer pH 7.4. 10 µl of fluorescent marker was mixed with 390 µl of TRIS buffer pH 7.4, and was used as a control to study the effect of organic solvents on fluorescence. Fluorescence of 10 µl of the final solutions along with 200 µl of TRIS buffer pH 7.4 was recorded with a fluorescent plate reader and results were compared.

**Figure B** represents the effects of extraction solvents on the fluorescence of the various markers. Tissue lysis buffer RIPA and organic solvents (methanol and acetonitrile) had minimal effects on the fluorescence of Na-Fl. On the other hand, for FITC-albumin samples prepared in RIPA buffer, methanol and acetonitrile, a quenching effect was experienced on the fluorescence of marker in the presence of these solvents. The quenching effect due to these organic solvents was reversed in samples prepared in solvent, dried and reconstituted in TRIS buffer pH 7.4. For FITC-dextran, samples prepared in acetonitrile experienced quenching effects due to organic solvent, which was reversed in samples prepared in solvent, dried and reconstituted in TRIS buffer pH 7.4 as indicated by enhanced fluorescence. Organic extracting solvent 1-butanol, enhanced the fluorescence of all three fluorescent markers illustrated by enhanced fluorescence of Na-Fl, FITC-albumin and FITC-dextran in aqueous phase, organic phase accompanied by samples prepared, dried and re-constituted in TRIS buffer.

To summarize the findings, Na-Fl fluorescence was stable in extracting solvents and methods, except in the process involving use of 1-butanol. For FITC-albumin and FITC-dextran, fluorescence was stable when fluorescent marker solutions were prepared in methanol and acetonitrile followed by drying and re-constitution in TRIS buffer. Additionally, FITC-dextran fluorescence was stable in samples prepared with lysis RIPA buffer or methanol only. Use of 1-butanol for the extraction and analysis procedures is therefore not advisable due to the enhancing effect of organic solvent over the fluorescence of Na-Fl, FITC-albumin and FITC-dextran.

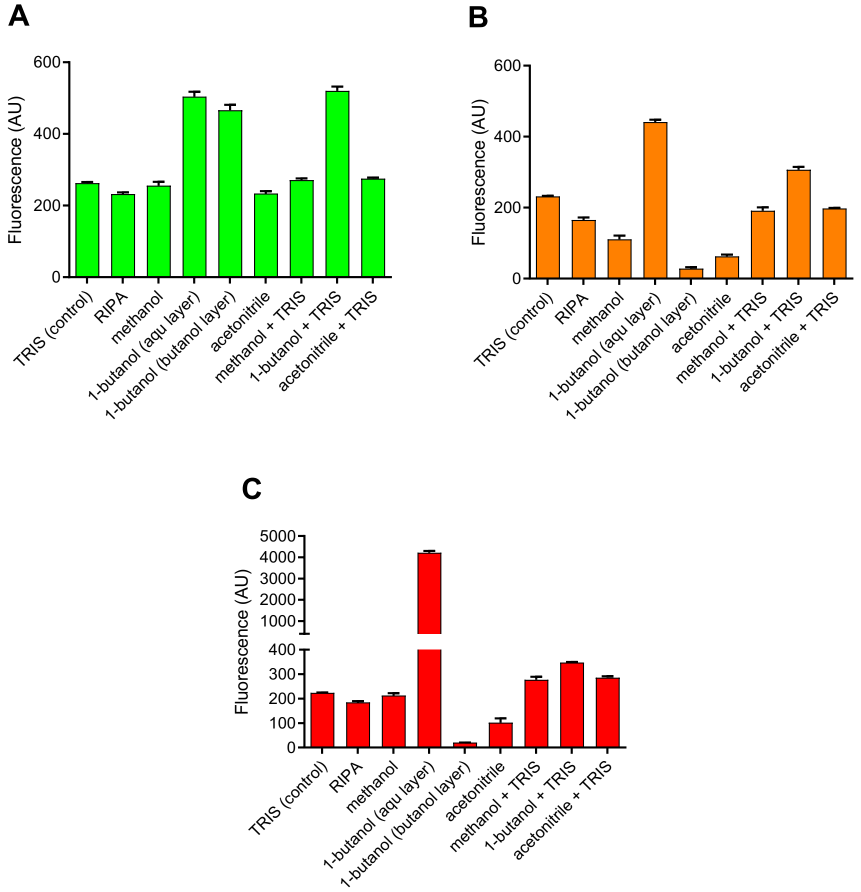


**Figure B: Effect of extracting solvent on fluorescence**: Bar graphs represent fluorescence level of sodium fluorescein (Na-Fl) (**A**), FITC-albumin (**B**) and FITC-dextran (**C**) in various organic solvents. Samples prepared and dried with organic solvent followed by re-constitution in TRIS buffer (pH 7.4). Results are compared with freshly prepared fluorescent marker solution's fluorescence prepared in TRIS buffer (pH 7.4) at the same volume. Data represents mean ± SEM of n=4 values.

*Extraction efficiency, extraction recovery and matrix effect:* Quantitative determination of fluorescent markers depends on two main factors: procedure efficiency as determined by extraction efficiency and extraction recovery, and secondly, interaction of markers with matrix as determined by the matrix effect. Hence, it was important to identify efficiency of extraction procedures and potential matrix interactions. This issue was solved by analyzing and comparing fluorescence of control samples (C) (10 µl of fluorescent marker i.e. 0.1% Na-Fl or 1% FITC-albumin or 1% FITC-dextran with 200 µl of TRIS buffer pH 7.4), samples spiked before extraction (BE) (i.e. 100 µl of brain homogenate spiked with either marker, incubated overnight, vortexed followed by extraction process using various solvents as explained in "extraction solvent effect" section), and samples spiked after extraction procedure (AE) (i.e. extraction of 100 µl of brain homogenate using various solvents as explained in "extraction solvent effect" section and spiking the resulting supernatant with either of fluorescent marker). Extraction efficiency, extraction recovery and matrix effects were calculated using following equations:

$$\textbf{Extraction efficiency} = (BE/C) * 100$$

$$\textbf{Extraction recovery} = (BE/AE) * 100$$

$$\textbf{Matrix effect} = (1- (AE/C)) * 100$$

Where **C** is control samples, i.e. standard solution without organic solvent treatment and without matrix; **BE** is before extraction samples, matrix sample spiked with marker before extraction; and **AE** is after extraction samples, matrix sample spiked with marker after extraction procedure. Positive matrix effect values reflects fluorescence enhancement, while negative matrix effect values reflect fluorescence suppression (quenching), due to matrix and solvent interaction with the fluorescent marker.

Extraction efficiency (EE) and extraction recovery (ER) of fluorescent markers from brain matrix using various extracting solvents is illustrated in **Figure C** and **Figure D**. Organic solvents resulted in better EE and ER of Na-Fl and FITC-albumin from matrix compared to TRIS buffer and tissue lysis RIPA buffer as illustrated. For Na-Fl, process EE and ER were 85% and 91% when extraction was performed by methanol only, and 92% and 89% when extraction of marker was performed by methanol and dried supernatant samples were reconstituted in TRIS buffer. Acetonitrile resulted in better extraction and recovery of Na-Fl from brain matrix in comparison to methanol, with values in a range of 82% and 94% respectively. However the process efficiency was improved when extraction of Na-Fl from brain matrix was performed by acetonitrile, and extracted supernatant dried and reconstituted in TRIS buffer at pH 7.4, resulting values in range of 99% of EE and 94% of ER.

For FITC-albumin, EE and ER values by methanol and acetonitrile extraction were low i.e. in range of 68% and 84% during extraction by methanol and 52% and 77% by acetonitrile extraction. Low values of FITC-albumin extractions by organic solvent could be quenching effect of solvents on the FITC-albumin fluorescence as explained in the previous section. Likewise Na-Fl, EE and ER of FITC-albumin was improved when extraction of marker was performed in organic solvents followed by drying of supernatant and reconstituted in TRIS buffer at pH 7.4. In comparison to acetonitrile extraction, methanol + TRIS buffer extraction yields improved EE and ER values i.e. 78% and 96% for FITC-albumin. Conversely, process efficiency of FITC-dextran was better in buffers when compared to organic solvents as indicated by EE and ER values in the range of 99% when extraction was performed by TRIS buffer at pH 7.4. EE and ER values were reduced (i.e. 93% and 83% respectively) when extraction was performed by RIPA tissue lysis buffer.

To summarize the results from Figure C and Figure D, for Na-Fl and FITC-albumin, better extraction of markers was achieved when extraction was performed by the acetonitrile + TRIS method for Na-Fl, methanol + TRIS for FITC-albumin, and TRIS extraction for FITC-dextran. Organic solvent 1-butanol is not a suitable extracting solvent as it provides false results by enhancing the fluorescence of markers, as also shown in the prior experiments.

**Figure E** illustrates the influence of matrix (i.e. brain homogenate) over the fluorescence of markers during the extraction procedure. The matrix effect reflects matrix-solvent interaction, matrix-marker interaction (responsible for extraction of marker) and marker-solvent interaction in the presence of matrix (responsible for fluorescence signal intensity change which affects quantitation of marker). Matrix interaction results in fluorescence suppression of Na-Fl when extraction was performed by methods involving methanol (-12%), acetonitrile (-9%), methanol + TRIS (-10%) and acetonitrile + TRIS (-4%). Likewise, Na-Fl, matrix interactions were responsible for fluorescence suppression when extraction was performed by methods involving methanol (-20%), acetonitrile (-22%), methanol + TRIS (-12%) and acetonitrile + TRIS (-15%). In the case of FITC-dextran, matrix produces mixed effect of fluorescence levels as indicated in the method involving organic solvents (i.e. methanol (+9%), acetonitrile (-98%), methanol + TRIS (-12%) and acetonitrile + TRIS (-15%)) and buffer solvents (i.e. 11% in TRIS buffer and 22% in RIPA buffer).

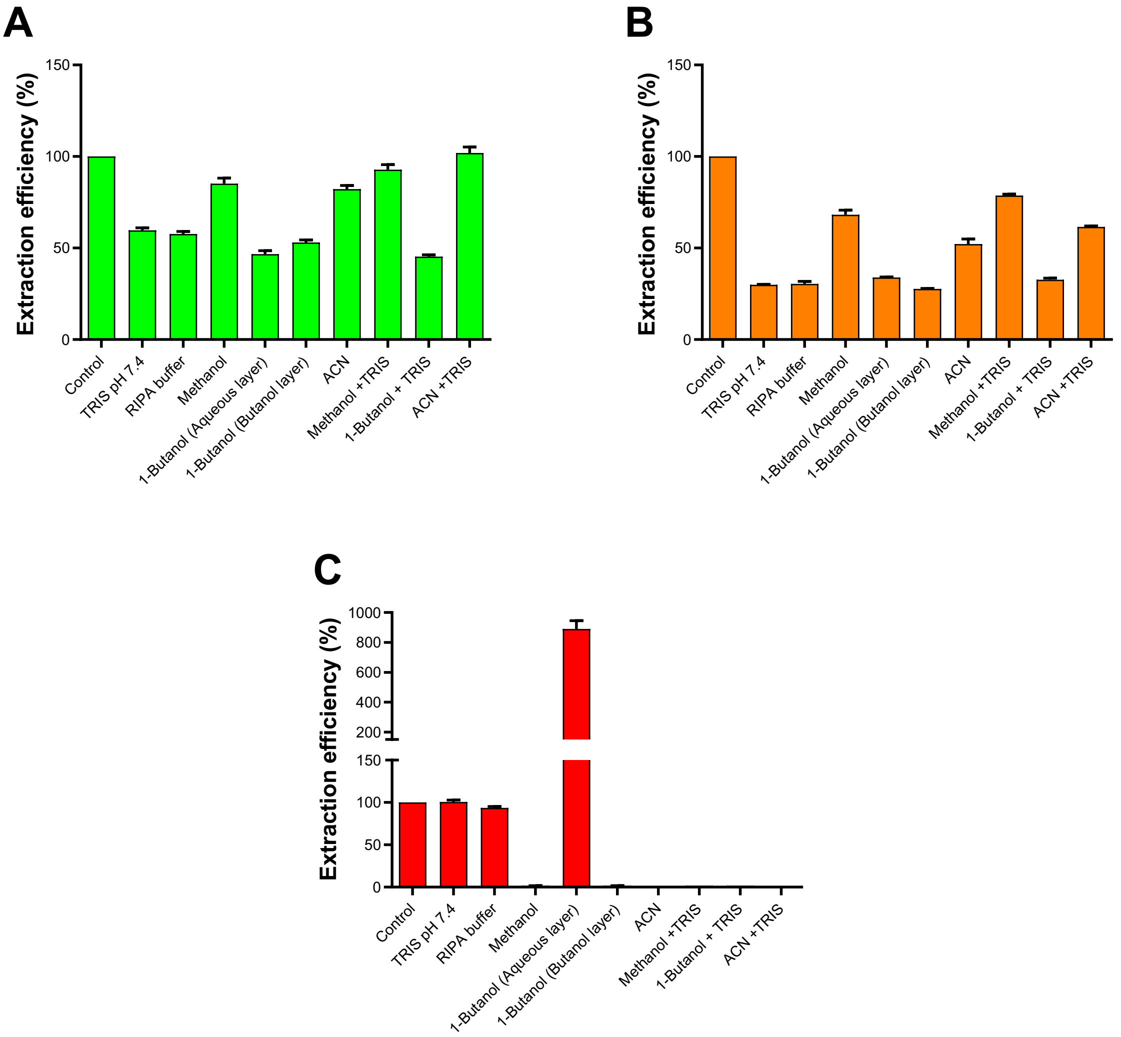


**Figure C: Extraction efficiency of process:** Bar graphs illustrates process efficiency as expressed by extraction efficiency of method used for the extraction of different fluorescent markers i.e. sodium fluorescein (**A**), FITC-albumin (**B**) and FITC-dextran (**C**) from brain homogenate matrix using different solvents. Results are compared with freshly prepared fluorescent marker solution's fluorescence prepared in TRIS buffer pH 7.4 at same volume in absence of matrix. Data represents mean ± SEM of n=4 values.

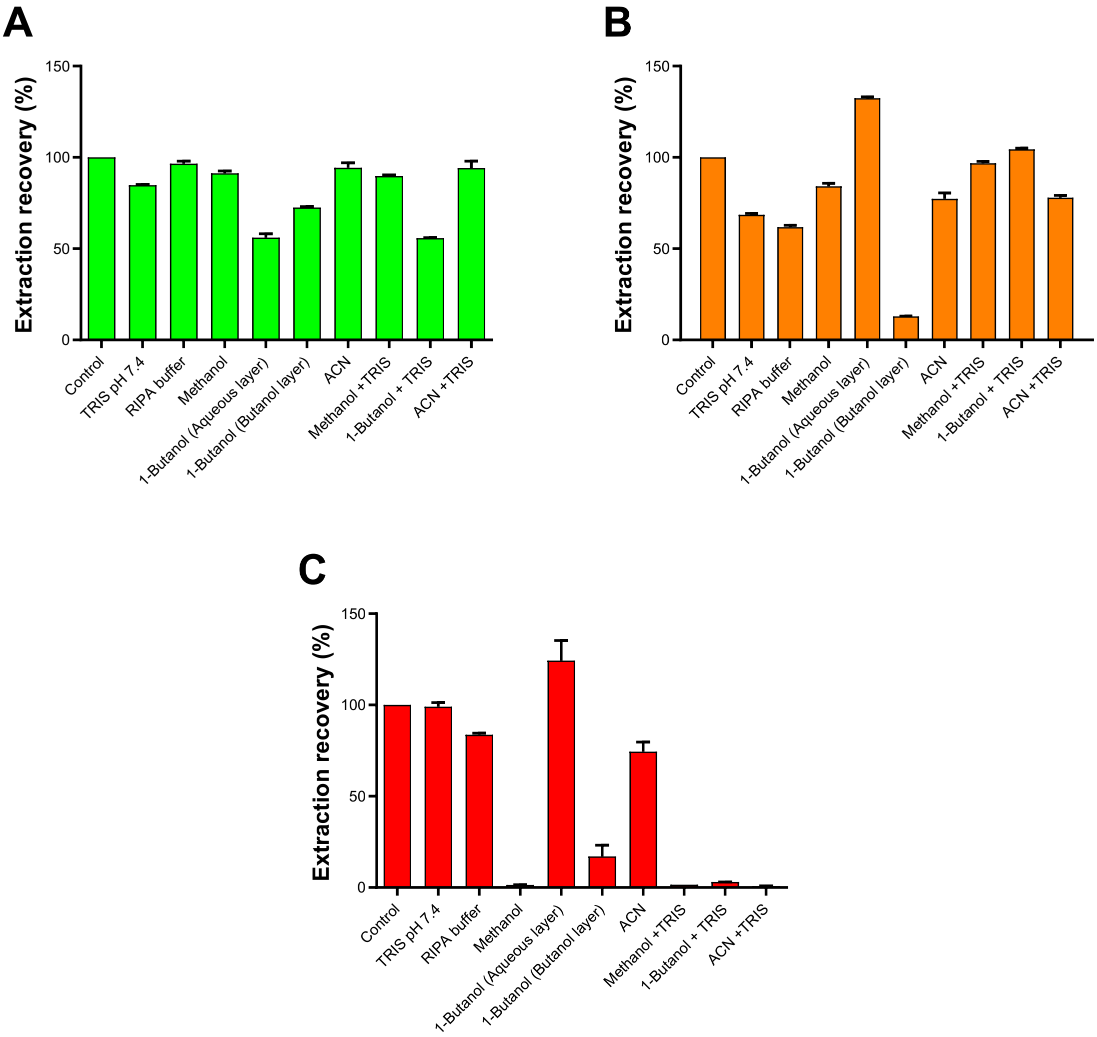


**Figure D: Extraction recovery of process:** Bar graphs illustrates process efficiency of extraction procedure as expressed by recovery of different fluorescent markers i.e. sodium fluorescein (**A**), FITC-albumin (**B**) and FITC-dextran (**C**) from brain homogenate matrix using different solvents. Results are compared with freshly prepared fluorescent marker solution's fluorescence prepared in TRIS buffer pH 7.4 at same volume in absence of matrix. Data represents mean ± SEM of n=4 values.

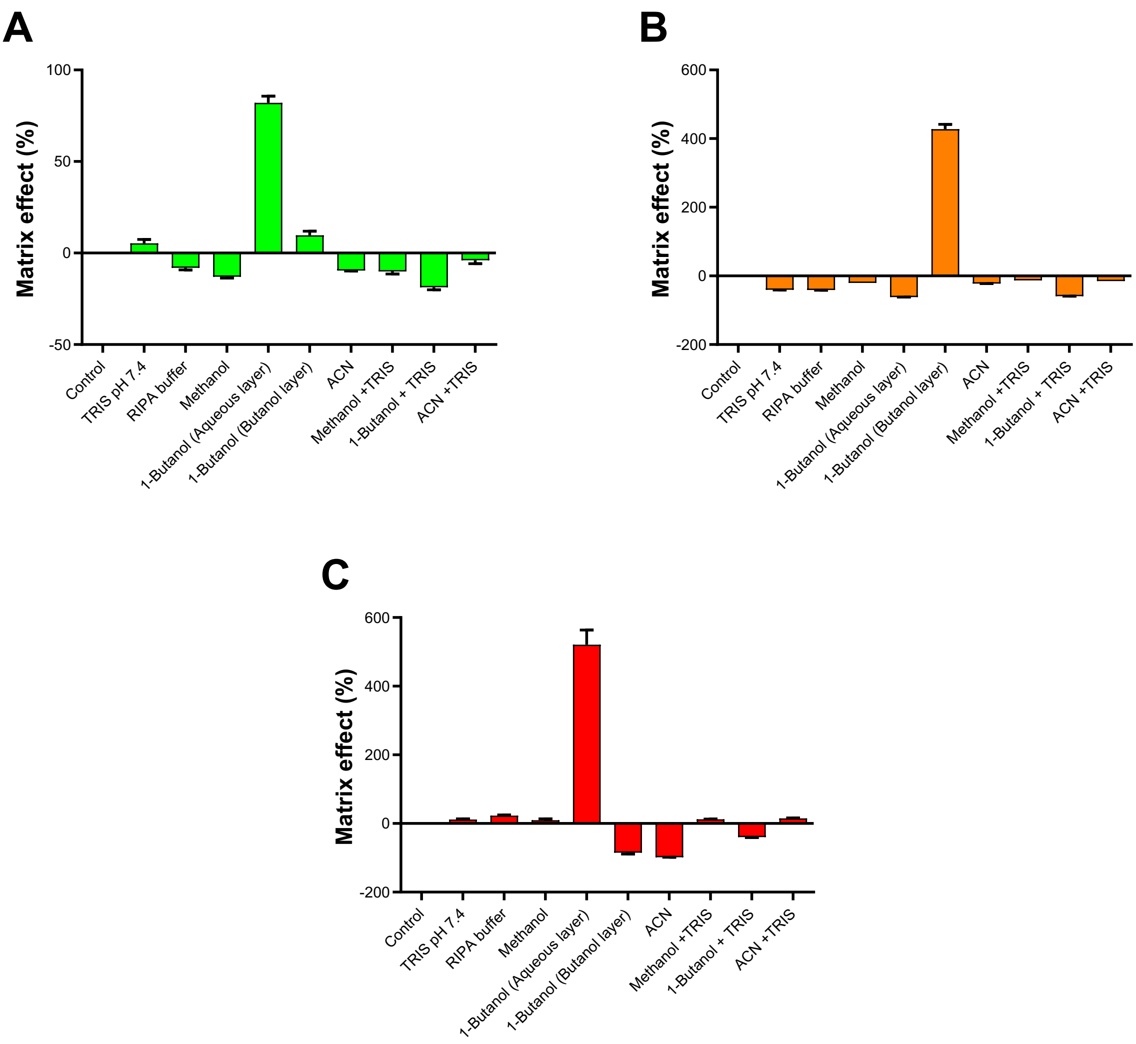


**Figure E: Influence of matrix and matrix interaction on fluorescence**: Bar graphs illustrates the influence of interaction between matrix (brain homogenate), solvents and fluorescent markers resulting in fluorescence enhancement (represented by positive values) or suppression (represented by negative values) of sodium fluorescein (**A**), FITC-albumin (**B**) and FITC-dextran (**C**). Results are compared with freshly prepared fluorescent marker solution's fluorescence prepared in TRIS buffer pH 7.4 at same volume in absence of matrix. Data represents mean ± SEM of n=4 values.

*Fluorescence loss:* Fluorescence loss of Na-Fl, FITC-albumin and FITC-dextran during extraction procedures was studied by re-suspending the precipitated pellet after extraction in NaOH solution. 100 µl of brain homogenates were spiked with 10 µl of either of fluorescent marker (i.e. 0.1% Na-Fl or 1% FITC-albumin or 1% FITC-dextran) and vortexed. Extraction of markers from spiked samples was then performed by protein precipitation using 300 µl of organic solvent (i.e. methanol or ACN or 1-butanol) followed by removal of supernatant. The residual pellet was incubated overnight in 400 µl of 5N NaOH solution. Results were compared to freshly prepared fluorescent marker solutions made by spiking 10 µl of same concentration in 390µl of TRIS buffer at pH 7.4. Fluorescence of 10µl of final solutions along with 200 µl of TRIS buffer as a loading solution was recorded using a fluorescent plate reader. Organic solvent used for extraction of fluorescent markers from tissue homogenates resulted in very minimal loss of the fluorescent marker (**Figure F**). Again however, in comparison to methanol and acetonitrile, 1-Butanol enhanced fluorescence resulting in increased fluorescence levels of Na-Fl, FITC-albumin and FITC-dextran.

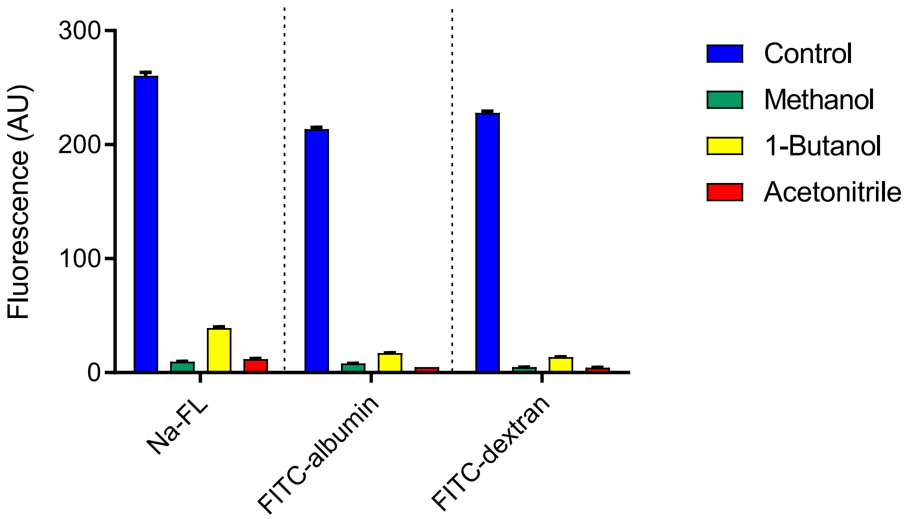


**Figure F: Fluorescence loss due to extraction procedure**: Extraction using organic solvent results in negligible 'loss' of fluorescent marker and fluorescence (< 0.1%). Bar graphs represents fluorescence level of sodium fluorescein (Na-Fl), FITC-albumin and FITC-dextran in supernatant of brain homogenate pellet obtained after protein precipitation and overnight incubation of pellet in NaOH solution. Results are compared with freshly prepared fluorescent marker solution's fluorescence prepared in TRIS buffer pH 7.4 at same volume. Data represents mean ± SEM of n=4 values.

In summary, the results from process stability, process efficiency and matrix effect experiments indicates that out of all solvents and extraction methods, acetonitrile + TRIS, methanol + TRIS, and TRIS alone, are suitable methods for Na-Fl, FITC-albumin and FITC-dextran extraction and recovery respectively. These selected methods resulted in minimal matrix effects and minimal interference for quantitative determination of fluorescent markers as summarized in the **Table** below.

**Table: Identification and selection of a favourable solvent system for quantification of fluorescent markers**

| Fluorescent Marker | Validation test | Advantageous solvent system & extraction method | | | | | | | |
|---|---|---|---|---|---|---|---|---|---|
| | | TRIS pH 7.4 | RIPA Buffer | Methanol | 1-Butanol | Acetonitrile | Methanol + TRIS pH 7.4 | 1-Butanol + TRIS pH 7.4 | Acetonitrile + TRIS pH 7.4 |
| Sodium Fluorescein | Loading solvent effect | | | | | | | | |
| | Extracting solvent effect | | | | | | | | |
| | Extraction efficiency | | | | | | | | |
| | Extraction recovery | | | | | | | | |
| | Matrix effect | | | | | | | | |
| | Fluorescence loss | | | | | | | | |
| FITC-albumin | Loading solvent effect | | | | | | | | |
| | Extracting solvent effect | | | | | | | | |
| | Extraction efficiency | | | | | | | | |
| | Extraction recovery | | | | | | | | |
| | Matrix effect | | | | | | | | |
| | Fluorescence loss | | | | | | | | |
| FITC-dextran | Loading solvent effect | | | | | | | | |
| | Extracting solvent effect | | | | | | | | |
| | Extraction efficiency | | | | | | | | |
| | Extraction recovery | | | | | | | | |
| | Matrix effect | | | | | | | | |
| | Fluorescence loss | | | | | | | | |

**Supplementary Data - all raw data used in publication**

**% uptake 15 min**

| | Extrapolated levels of marker (ng) | | | | | | | | | | | | | | |
|---|---|---|---|---|---|---|---|---|---|---|---|---|---|---|---|
| | Serum (10µl) | | | | | Brain (10mg) | | | | | %uptake | | | | |
| NaFl_Saline | 33800.05 | 32633.63 | 30854.39 | 37496.58 | 43735.98 | 2061.80 | 3934.59 | 2575.27 | 2081.36 | 4661.29 | 6.10 | 12.06 | 8.35 | 5.55 | 10.66 |
| NaFl_LPS | 32669.75 | 32416.69 | 21373.48 | 42467.68 | 29394.09 | 9545.77 | 11935.78 | 11728.07 | 16570.28 | 11560.85 | 29.22 | 36.82 | 54.87 | 39.02 | 39.33 |
| FITC-albumin_Saline | 32158.98 | 32168.20 | 31212.03 | 31988.30 | 38755.15 | 607.74 | 384.52 | 305.26 | 433.24 | 524.89 | 1.89 | 1.20 | 0.98 | 1.35 | 1.35 |
| FITC-albumin_LPS | 28037.93 | 33075.51 | 36733.50 | 404512.40 | 35159.25 | 1325.47 | 1663.80 | 1526.25 | 18010.40 | 2069.81 | 4.73 | 5.03 | 4.15 | 4.45 | 5.89 |

**% uptake 1 hr**

| | Extrapolated levels of marker (ng) | | | | | | | | | | | | | | |
|---|---|---|---|---|---|---|---|---|---|---|---|---|---|---|---|
| | Serum (10µl) | | | | | Brain (10mg) | | | | | %uptake | | | | |
| NaFl | 27314.41 | 25451.17 | 24891.89 | 30869.93 | 27964.47 | 15104.98 | 14548.24 | 13870.32 | 12606.80 | 11345.71 | 55.30 | 57.16 | 55.72 | 40.84 | 40.57 |
| FITC-albumin | 26280.84 | 49598.77 | 41556.02 | 24135.46 | 31598.72 | 2901.53 | 4162.95 | 11681.88 | 5248.70 | 4536.31 | 11.04 | 8.39 | 28.11 | 21.75 | 14.36 |
| FITC-dextran | 7397.93 | 7668.89 | 6379.73 | 7470.42 | 7735.81 | 120.22 | 66.41 | 86.76 | 46.24 | 138.20 | 1.63 | 0.87 | 1.36 | 0.62 | 1.79 |

**% uptake 24 hr**

| | Extrapolated levels of marker (ng) | | | | | | | | | | | | | | |
|---|---|---|---|---|---|---|---|---|---|---|---|---|---|---|---|
| | Serum (10µl) | | | | | Brain (10mg) | | | | | %uptake | | | | |
| NaFl_Saline | 10447.12 | 11382.41 | 13114.69 | 12887.26 | 17741.97 | 357.71 | 371.04 | 428.25 | 573.74 | 702.19 | 3.42 | 3.26 | 3.27 | 4.45 | 3.96 |
| NaFl_LPS | 9771.16 | 10386.18 | 11504.73 | 10971.91 | 12225.56 | 1686.39 | 2453.57 | 2543.52 | 1328.43 | 1354.12 | 17.26 | 23.62 | 22.11 | 12.11 | 11.08 |
| FITC-albumin_Saline | 17749.96 | 14951.08 | 14192.11 | 13629.14 | 14684.58 | 1959.68 | 857.84 | 1191.18 | 1144.21 | 1234.97 | 11.04 | 5.74 | 8.39 | 8.40 | 8.41 |
| FITC-albumin_LPS | 12326.55 | 12168.80 | 13307.55 | 15159.25 | 13673.35 | 2913.64 | 3060.65 | 2764.59 | 3374.73 | 4024.72 | 23.64 | 25.15 | 20.77 | 22.26 | 29.43 |

**% uptake WT vs SOD1**

*Spinal cord*

| | WT | | | | | SOD1 | | | | |
|---|---|---|---|---|---|---|---|---|---|---|
| P30 | 7.39 | 7.84 | 8.13 | 7.60 | 7.05 | 9.47 | 6.61 | 7.78 | 7.96 | 7.96 |
| P70 | 10.18 | 9.88 | 11.20 | 10.04 | 8.89 | 10.98 | 11.64 | 12.08 | 11.30 | 10.48 |
| P130 | 11.40 | 10.17 | 10.26 | 10.25 | 9.16 | 12.47 | 15.16 | 12.19 | 13.25 | 13.19 |
| P175 | 14.28 | 14.10 | 12.85 | 13.48 | 12.69 | 28.50 | 27.40 | 24.41 | 26.33 | 25.02 |

*Cortex*

| | WT | | | | | SOD1 | | | | |
|---|---|---|---|---|---|---|---|---|---|---|
| P30 | 4.77 | 4.48 | 5.24 | 4.63 | 4.03 | 4.82 | 5.33 | 5.80 | 5.31 | 5.31 |
| P70 | 5.46 | 5.92 | 6.01 | 5.68 | 5.32 | 5.94 | 7.78 | 6.53 | 6.81 | 7.00 |
| P130 | 7.15 | 7.03 | 7.87 | 7.09 | 6.33 | 10.18 | 10.55 | 11.28 | 10.35 | 9.37 |
| P175 | 9.33 | 8.81 | 10.27 | 9.09 | 7.93 | 14.62 | 15.23 | 14.25 | 14.21 | 12.75 |

*Striatum*

| | WT | | | | | SOD1 | | | | |
|---|---|---|---|---|---|---|---|---|---|---|
| P30 | 3.61 | 6.11 | 3.97 | 4.80 | 5.50 | 4.11 | 6.99 | 4.53 | 5.48 | 6.29 |
| P70 | 6.46 | 7.55 | 7.11 | 6.98 | 6.80 | 9.89 | 9.73 | 9.01 | 9.54 | 9.54 |
| P130 | 7.96 | 7.49 | 7.53 | 7.43 | 6.74 | 11.58 | 10.75 | 10.68 | 11.00 | 11.00 |
| P175 | 8.52 | 8.35 | 9.37 | 8.44 | 7.51 | 12.99 | 12.05 | 11.63 | 12.16 | 11.96 |

*Cerebellum*

| | WT | | | | | SOD1 | | | | |
|---|---|---|---|---|---|---|---|---|---|---|
| P30 | 7.81 | 8.28 | 8.59 | 8.03 | 7.45 | 8.81 | 6.99 | 9.69 | 7.94 | 6.29 |
| P70 | 10.40 | 10.09 | 11.45 | 10.26 | 9.08 | 12.75 | 11.90 | 10.47 | 11.71 | 11.71 |
| P130 | 12.14 | 10.83 | 13.35 | 11.52 | 9.75 | 13.28 | 16.14 | 12.98 | 14.11 | 14.04 |
| P175 | 10.79 | 10.65 | 11.86 | 10.72 | 9.58 | 21.53 | 20.70 | 18.44 | 19.89 | 18.90 |

**% uptake WT vs R6/1**

*Spinal cord*

| | WT | | | | | R6/1 | | | | |
|---|---|---|---|---|---|---|---|---|---|---|
| 6 weeks | 10.76 | 6.81 | 14.91 | 3.91 | 10.82 | 11.02 | 7.14 | 16.07 | 4.25 | 9.62 |
| 12 weeks | 12.44 | 7.49 | 18.02 | 4.70 | 12.65 | 18.47 | 13.86 | 21.23 | 11.16 | 16.18 |
| 20 weeks | 17.84 | 14.77 | 5.62 | 3.95 | 12.74 | 50.38 | 20.83 | 22.71 | 22.85 | 29.19 |
| 32 weeks | 20.98 | 5.57 | 19.20 | 5.85 | 15.25 | 38.91 | 39.96 | 33.17 | 31.00 | 35.76 |

*Cortex*

| | WT | | | | | R6/1 | | | | |
|---|---|---|---|---|---|---|---|---|---|---|
| 6 weeks | 3.04 | 1.70 | 5.15 | 6.82 | 3.30 | 3.05 | 1.94 | 6.32 | 7.21 | 4.63 |
| 12 weeks | 4.30 | 2.55 | 8.30 | 9.50 | 5.05 | 10.41 | 9.90 | 12.02 | 7.25 | 9.89 |
| 20 weeks | 6.32 | 5.20 | 5.55 | 4.95 | 5.69 | 10.30 | 10.70 | 11.65 | 14.11 | 11.69 |
| 32 weeks | 10.76 | 5.91 | 7.80 | 4.37 | 8.16 | 15.29 | 21.30 | 15.43 | 16.66 | 17.17 |

*Striatum*

| | WT | | | | | R6/1 | | | | |
|---|---|---|---|---|---|---|---|---|---|---|
| 6 weeks | 2.80 | 1.54 | 7.11 | 6.06 | 3.82 | 2.86 | 1.89 | 8.22 | 6.14 | 4.78 |
| 12 weeks | 3.16 | 1.96 | 8.86 | 6.78 | 4.66 | 11.92 | 10.03 | 10.37 | 4.94 | 9.31 |
| 20 weeks | 7.05 | 4.40 | 6.37 | 4.98 | 5.94 | 19.78 | 9.97 | 13.75 | 12.54 | 14.01 |
| 32 weeks | 8.54 | 6.85 | 8.29 | 3.58 | 7.89 | 21.31 | 22.09 | 17.40 | 20.14 | 20.23 |

*Cerebellum*

| | WT | | | | | R6/1 | | | | |
|---|---|---|---|---|---|---|---|---|---|---|
| 6 weeks | 4.61 | 2.12 | 12.54 | 13.30 | 6.42 | 4.31 | 2.66 | 14.89 | 14.51 | 9.09 |
| 12 weeks | 5.71 | 2.85 | 15.18 | 16.58 | 7.91 | 13.41 | 10.91 | 9.58 | 8.81 | 10.68 |
| 20 weeks | 8.35 | 5.26 | 8.93 | 17.58 | 7.51 | 19.86 | 14.45 | 10.13 | 9.96 | 13.60 |
| 32 weeks | 17.98 | 8.06 | 21.50 | 7.62 | 15.84 | 31.72 | 26.46 | 28.29 | 26.87 | 28.34 |

**% uptake WT vs APP/PS1**

*Spinal cord*

| | WT | | | | | APP/PS1 | | | | |
|---|---|---|---|---|---|---|---|---|---|---|
| 4 months | 8.89 | 3.70 | 8.46 | 10.15 | 6.87 | 10.27 | 10.70 | 10.73 | 10.30 | 11.04 |
| 18 months | 7.88 | 11.89 | 10.81 | 11.12 | 8.45 | 6.30 | 5.46 | 4.67 | 7.03 | 3.23 |

*Cortex*

| | WT | | | | | APP/PS1 | | | | |
|---|---|---|---|---|---|---|---|---|---|---|
| 4 months | 4.68 | 5.67 | 3.63 | 5.12 | 4.02 | 6.71 | 4.87 | 7.39 | 4.93 | 5.08 |
| 18 months | 6.11 | 6.67 | 3.37 | 5.58 | 6.72 | 4.05 | 3.57 | 4.09 | 4.85 | 4.21 |

*Striatum*

| | WT | | | | | APP/PS1 | | | | |
|---|---|---|---|---|---|---|---|---|---|---|
| 4 months | 4.82 | 4.70 | 5.15 | 4.69 | 4.94 | 6.91 | 6.39 | 8.39 | 7.41 | 6.84 |
| 18 months | 10.65 | 11.47 | 5.23 | 4.40 | 3.27 | 3.76 | 3.82 | 2.68 | 4.69 | 4.04 |

*Cerebellum*

| | WT | | | | | APP/PS1 | | | | |
|---|---|---|---|---|---|---|---|---|---|---|
| 4 months | 9.74 | 6.47 | 7.47 | 9.07 | 7.46 | 8.91 | 12.89 | 12.27 | 7.91 | 9.61 |
| 18 months | 12.37 | 7.58 | 13.45 | 8.30 | 9.55 | 6.93 | 7.54 | 6.27 | 10.65 | 6.31 |

*Hippocampus*

| | WT | | | | | APP/PS1 | | | | |
|---|---|---|---|---|---|---|---|---|---|---|
| 4 months | 7.83 | 9.87 | 5.74 | 8.62 | 8.64 | 9.73 | 10.82 | 7.25 | 10.53 | 8.43 |
| 18 months | 8.59 | 9.80 | 13.03 | 9.80 | 9.18 | 6.88 | 3.68 | 2.96 | 7.11 | 5.80 |

**NaFl amount WT vs SOD1**

*Spinal cord*

| | WT | | | | | SOD1 | | | | |
|---|---|---|---|---|---|---|---|---|---|---|
| P30 | 2.34 | 2.47 | 2.55 | 2.37 | 2.18 | 2.91 | 1.98 | 2.30 | 2.35 | 2.31 |
| P70 | 2.90 | 2.78 | 3.10 | 2.70 | 2.35 | 2.83 | 2.92 | 2.99 | 2.79 | 2.58 |
| P130 | 2.90 | 2.53 | 3.37 | 3.34 | 2.98 | 4.06 | 4.87 | 3.91 | 4.25 | 4.22 |
| P175 | 4.52 | 4.45 | 4.01 | 4.20 | 3.95 | 8.82 | 8.45 | 7.52 | 8.10 | 7.66 |

*Cortex*

| | WT | | | | | SOD1 | | | | |
|---|---|---|---|---|---|---|---|---|---|---|
| P30 | 1.51 | 1.41 | 1.64 | 1.44 | 1.24 | 1.48 | 1.60 | 1.71 | 1.57 | 1.54 |
| P70 | 1.56 | 1.66 | 1.66 | 1.53 | 1.41 | 1.53 | 1.95 | 1.62 | 1.68 | 1.72 |
| P130 | 1.82 | 1.75 | 2.58 | 2.31 | 2.06 | 3.31 | 3.39 | 3.62 | 3.32 | 3.00 |
| P175 | 2.96 | 2.78 | 3.20 | 2.83 | 2.47 | 4.53 | 4.70 | 4.39 | 4.37 | 3.90 |

*Striatum*

| | WT | | | | | SOD1 | | | | |
|---|---|---|---|---|---|---|---|---|---|---|
| P30 | 1.14 | 1.93 | 1.24 | 1.50 | 1.70 | 1.27 | 2.09 | 1.34 | 1.62 | 1.83 |
| P70 | 1.84 | 2.12 | 1.97 | 1.88 | 1.80 | 2.55 | 2.44 | 2.23 | 2.36 | 2.35 |
| P130 | 2.02 | 1.86 | 2.47 | 2.42 | 2.19 | 3.77 | 3.45 | 3.43 | 3.53 | 3.52 |
| P175 | 2.70 | 2.63 | 2.92 | 2.63 | 2.34 | 4.02 | 3.71 | 3.58 | 3.74 | 3.66 |

*Cerebellum*

| | WT | | | | | SOD1 | | | | |
|---|---|---|---|---|---|---|---|---|---|---|
| P30 | 2.47 | 2.61 | 2.69 | 2.50 | 2.30 | 2.71 | 2.09 | 2.86 | 2.34 | 1.83 |
| P70 | 2.97 | 2.84 | 3.16 | 2.76 | 2.41 | 3.29 | 2.98 | 2.59 | 2.89 | 2.88 |
| P130 | 3.09 | 2.69 | 4.38 | 3.75 | 3.17 | 4.32 | 5.18 | 4.16 | 4.52 | 4.50 |
| P175 | 3.42 | 3.36 | 3.70 | 3.34 | 2.98 | 6.66 | 6.38 | 5.68 | 6.12 | 5.79 |

*Serum*

| | WT | | | | | SOD1 | | | | |
|---|---|---|---|---|---|---|---|---|---|---|
| P30 | 31.68 | 31.55 | 31.34 | 31.18 | 30.83 | 30.78 | 29.96 | 29.55 | 29.50 | 29.03 |
| P70 | 28.51 | 28.14 | 27.65 | 26.95 | 26.48 | 25.79 | 25.06 | 24.75 | 24.72 | 24.58 |
| P130 | 25.42 | 24.87 | 32.81 | 32.57 | 32.56 | 32.55 | 32.11 | 32.10 | 32.05 | 32.03 |
| P175 | 31.68 | 31.55 | 31.18 | 31.14 | 31.11 | 30.94 | 30.83 | 30.80 | 30.77 | 30.61 |

**NaFl amount WT vs R6/1**

*Spinal cord*

| | WT | | | | | R6/1 | | | | |
|---|---|---|---|---|---|---|---|---|---|---|
| 6 weeks | 3.39 | 2.15 | 4.68 | 1.22 | 3.36 | 3.42 | 2.19 | 4.78 | 1.21 | 2.71 |
| 12 weeks | 3.46 | 2.07 | 4.92 | 1.28 | 3.40 | 4.91 | 3.62 | 5.21 | 2.74 | 4.01 |
| 20 weeks | 4.60 | 3.80 | 1.44 | 1.01 | 3.21 | 12.69 | 5.23 | 5.67 | 7.47 | 9.41 |
| 32 weeks | 6.76 | 1.79 | 6.14 | 1.87 | 4.83 | 12.19 | 12.32 | 10.19 | 9.53 | 10.90 |

*Cortex*

| | WT | | | | | R6/1 | | | | |
|---|---|---|---|---|---|---|---|---|---|---|
| 6 weeks | 0.96 | 0.54 | 1.62 | 2.13 | 1.03 | 0.95 | 0.60 | 1.88 | 2.05 | 1.30 |
| 12 weeks | 1.20 | 0.70 | 2.27 | 2.59 | 1.36 | 2.77 | 2.59 | 2.95 | 1.78 | 2.45 |
| 20 weeks | 1.63 | 1.34 | 1.42 | 1.26 | 1.43 | 2.59 | 2.69 | 2.91 | 4.61 | 3.77 |
| 32 weeks | 3.47 | 1.89 | 2.50 | 1.40 | 2.59 | 4.79 | 6.57 | 4.74 | 5.12 | 5.23 |

*Striatum*

| | WT | | | | | R6/1 | | | | |
|---|---|---|---|---|---|---|---|---|---|---|
| 6 weeks | 0.88 | 0.49 | 2.23 | 1.90 | 1.19 | 0.89 | 0.58 | 2.44 | 1.75 | 1.34 |
| 12 weeks | 0.88 | 0.54 | 2.42 | 1.85 | 1.25 | 3.17 | 2.62 | 2.55 | 1.21 | 2.31 |
| 20 weeks | 1.82 | 1.13 | 1.63 | 1.27 | 1.50 | 4.98 | 2.51 | 3.43 | 4.10 | 4.52 |
| 32 weeks | 2.75 | 2.20 | 2.65 | 1.14 | 2.50 | 6.67 | 6.81 | 5.35 | 6.19 | 6.16 |

*Cerebellum*

| | WT | | | | | R6/1 | | | | |
|---|---|---|---|---|---|---|---|---|---|---|
| 6 weeks | 1.46 | 0.67 | 3.93 | 4.16 | 2.00 | 1.34 | 0.81 | 4.43 | 4.13 | 2.56 |
| 12 weeks | 1.59 | 0.79 | 4.14 | 4.52 | 2.13 | 3.57 | 2.85 | 2.35 | 2.16 | 2.65 |
| 20 weeks | 2.15 | 1.35 | 2.28 | 4.47 | 1.89 | 5.00 | 3.63 | 2.53 | 3.25 | 4.39 |
| 32 weeks | 5.79 | 2.58 | 6.88 | 2.43 | 5.02 | 9.93 | 8.16 | 8.69 | 8.26 | 8.63 |

*Serum*

| | WT | | | | | R6/1 | | | | |
|---|---|---|---|---|---|---|---|---|---|---|
| 6 weeks | 31.54 | 31.52 | 31.36 | 31.31 | 31.08 | 31.01 | 30.63 | 29.75 | 28.44 | 28.15 |
| 12 weeks | 27.82 | 27.64 | 27.30 | 27.25 | 26.87 | 26.60 | 26.13 | 24.55 | 24.53 | 24.81 |
| 20 weeks | 25.79 | 25.72 | 25.55 | 25.45 | 25.19 | 25.18 | 25.13 | 24.96 | 32.69 | 32.25 |
| 32 weeks | 32.21 | 32.07 | 31.99 | 31.93 | 31.69 | 31.32 | 30.83 | 30.73 | 30.73 | 30.47 |

**NaFl amount WT vs APP/PS1**

| | WT | | | | | APP/PS1 | | | | |
|---|---|---|---|---|---|---|---|---|---|---|
| ***Spinal cord*** | | | | | | | | | | |
| 4 months | 2.84 | 1.17 | 2.63 | 3.13 | 2.02 | 3.01 | 3.10 | 3.10 | 2.93 | 3.12 |
| 18 months | 2.19 | 3.24 | 2.87 | 2.95 | 2.16 | 1.56 | 1.34 | 1.16 | 2.12 | 0.97 |
| ***Cortex*** | | | | | | | | | | |
| 4 months | 1.49 | 1.79 | 1.13 | 1.58 | 1.18 | 1.97 | 1.41 | 2.14 | 1.40 | 1.44 |
| 18 months | 1.70 | 1.82 | 0.90 | 1.48 | 1.72 | 1.00 | 0.88 | 1.01 | 1.46 | 1.26 |
| ***Striatum*** | | | | | | | | | | |
| 4 months | 1.54 | 1.48 | 1.60 | 1.44 | 1.45 | 2.03 | 1.85 | 2.43 | 2.11 | 1.93 |
| 18 months | 2.97 | 3.13 | 1.39 | 1.17 | 0.84 | 0.93 | 0.94 | 0.66 | 1.41 | 1.21 |
| ***Cerebellum*** | | | | | | | | | | |
| 4 months | 3.11 | 2.04 | 2.32 | 2.79 | 2.20 | 2.61 | 3.74 | 3.55 | 2.25 | 2.71 |
| 18 months | 3.44 | 2.07 | 3.58 | 2.20 | 2.44 | 1.71 | 1.85 | 1.55 | 3.20 | 1.89 |
| ***Hippocampus*** | | | | | | | | | | |
| 4 months | 2.50 | 3.11 | 1.78 | 2.65 | 2.54 | 2.86 | 3.14 | 2.10 | 2.99 | 2.38 |
| 18 months | 2.39 | 2.67 | 3.47 | 2.60 | 2.34 | 1.70 | 0.91 | 0.73 | 2.14 | 1.74 |
| ***Serum*** | | | | | | | | | | |
| 4 months | 31.89 | 31.48 | 31.02 | 30.80 | 29.45 | 29.34 | 29.00 | 28.94 | 28.41 | 28.25 |
| 18 months | 27.84 | 27.27 | 26.59 | 26.50 | 25.55 | 24.70 | 24.58 | 24.78 | 30.09 | 30.00 |
